# A genome-guided multilocus sequence analysis scheme to detect and resolve diversity within the X-disease phytoplasma group (16SrIII)

**DOI:** 10.64898/2026.11.13.766501

**Authors:** Franco D. Fernández, Vanina A. Bongiorno, Luis R. Conci

## Abstract

Phytoplasmas of the X-disease group (16SrIII) cause economically important diseases of vegetables, forage and staple crops, fruit trees and ornamentals in South America, where they are diagnosed and classified only by 16S rRNA gene RFLP. Because this gene is highly conserved, strains that differ in host range, vectors and regulatory relevance cannot be distinguished. We developed a multilocus sequence analysis (MLSA) scheme for typing 16SrIII phytoplasmas, using genome data to select the markers and to define reference lineages. Five single-copy loci (*tuf*, *secA*, *secY*, *mutM*, *cimH*) were chosen from 16 curated genomes; primers matched all genomes *in silico*, and four of the five pairs did not amplify phytoplasmas of other groups co-occurring in the region (16SrI, 16SrVII, 16SrXIII). Genome comparisons resolved four lineages, including one restricted to South America that shared only 90.8–93.9% average nucleotide identity with the other lineages, despite > 98.8% 16S rRNA identity. The concatenated MLSA (2,870 nt) reproduced the genome-based lineages and assigned five Argentinian field isolates from weeds, fodder beet, cassava, plum and a native shrub. Identical 16SrIII-B RFLP patterns occurred in different lineages, and one 16SrIII-B isolate grouped within the 16SrIII-J clade, showing that subgroups are not reliable proxies for lineages. The South American lineage had the broadest host range, spanning six plant families. We also detected chimeric rRNA copies and sequencing artefacts that can cause misclassification. This MLSA provides a PCR-based tool to identify 16SrIII lineages in crops, weeds and insects, supporting epidemiological studies, disease management and phytosanitary decisions.

## Introduction

Phytoplasmas are wall-less, phloem-restricted bacteria of the class *Mollicutes* that cause yellowing, witches’ broom, phyllody, virescence, stunting and decline in hundreds of plant species worldwide, with major losses in crops and fruit trees (Bertaccini, 2022), and are transmitted by phloem-feeding insects, mainly leafhoppers and psyllids. Because they cannot be cultured, their detection and classification rely on molecular markers, primarily the 16S rRNA gene: groups and subgroups are defined by virtual RFLP of the R16F2n/R16R2 fragment (Zhao et al., 2009), and ‘*Candidatus* Phytoplasma’ species are delimited by 16S rRNA identity and, increasingly, by genome-based criteria such as average nucleotide identity (ANI; Bertaccini et al., 2022; Kirdat et al., 2023).

The X-disease group (16SrIII) includes ‘*Ca.* P. pruni’, the agent of X-disease of peach and cherry (Davis et al., 2013), which has caused recurrent epidemics and severe losses in North American stone fruit production; recent multilocus studies in that region revealed several lineages that differ in distribution and in the symptoms they induce (Molnar et al., 2024). In South America, where three decades of research have documented 16 ‘*Ca.* Phytoplasma’ species, ‘*Ca.* P. pruni’-related strains are among the most frequently detected phytoplasmas in agriculturally relevant crops and have been reported in Argentina, Brazil, Chile and Colombia, among other countries (Montano et al., 2024). In Argentina, the group is one of the most widespread: subgroups 16SrIII-B and 16SrIII-J predominate, and the new subgroups 16SrIII-W and 16SrIII-X were described from local hosts (Galdeano et al., 2013). These phytoplasmas are associated with diseases of vegetables (lettuce, garlic, Swiss chard), industrial, staple and forage crops (sunflower, cassava, fodder beet), fruit crops (peach, plum, cherry, grapevine, strawberry) and ornamental and forest trees such as chinaberry, while weeds and native plants act as natural reservoirs (Galdeano et al., 2013; Fernández et al., 2018, 2020; Bongiorno et al., 2020; Montano et al., 2024). Some of these diseases cause direct economic losses; for example, fodder beet wilting-yellowing, associated with 16SrIII-J, has emerged as a significant constraint on forage production for cattle in northern Patagonia and southern Buenos Aires province (Bongiorno et al., 2025).

The known diversity of the group in the region continues to expand. Recent work has documented new hosts and diseases, including cassava witches’ broom in Argentina and cassava oversprouting in Brazil (Fernández et al., 2018; Oliveira et al., 2026), plum yellows in mixed infections with ‘*Ca.* P. meliae’ (Bongiorno et al., 2020), and the spread of 16SrIII-X in lettuce crops (Fernández et al., 2026). Genome sequences are now available for South American strains of subgroups 16SrIII-B, -J and -X (Zamorano and Fiore, 2016; Fernández et al., 2020c, 2024, 2026; Oliveira et al., 2026). Strains of 16SrIII-J are transmitted by the leafhoppers *Paratanus exitiosus* and *Bergallia valdiviana* in Chile (Quiroga et al., 2019), and *P. exitiosus* was recently shown to transmit the fodder beet wilting-yellowing phytoplasma (16SrIII-J) in Argentina (Bongiorno et al., 2025); however, the vectors of most other South American strains remain unconfirmed (Montano et al., 2024), and markers distinguish North and South American lineages within the same 16Sr subgroups (Pérez-López et al., 2017). This combination of broad host range, uncharacterised vectors and regional lineages hampers epidemiological studies and management.

These questions also have regulatory consequences. ‘*Ca.* P. pruni’ (Western X disease) is included in the EPPO A1 list of pests recommended for regulation as quarantine pests (EPPO, 2024), but South American 16SrIII strains are usually reported as ‘*Ca.* P. pruni’-related on the basis of 16S rRNA data alone, and a distinct taxon, ‘*Ca.* P. platensis’, was proposed for 16SrIII-J strains on multilocus and biological grounds (Fernández et al., 2019). Whether strains detected in a crop belong to the same lineage as those causing X-disease elsewhere, or to regional lineages with different hosts and vectors, cannot be resolved with current diagnostic tools.

The 16S rRNA gene is too conserved to resolve this diversity: identical RFLP patterns can mask genomic divergence, the two rRNA operons may differ, and subgroup assignment does not necessarily reflect relationships elsewhere in the genome. Less conserved genes improve resolution: *secY* and ribosomal protein genes have been used to characterise ‘*Ca.* P. pruni’ and other groups (Lee et al., 2010; Davis et al., 2013), and the membrane protein genes *imp* and *idpA* separated the Italian 16SrIII-B strain MA1 from Argentinian 16SrIII-B strains (Alessio et al., 2025). However, multilocus markers have usually been chosen *a priori*, and their ability to reproduce genome-wide relationships has rarely been tested; for 16SrI phytoplasmas, genome data have been used to select and evaluate such markers (Cho et al., 2020), but no equivalent, field-applicable typing system exists for the X-disease group.

The aim of this study was to develop and validate a genome-guided multilocus sequence analysis (MLSA) scheme for typing X-disease group phytoplasmas. Specifically, we (1) selected single-copy loci from genome-wide orthology and designed primers evaluated in silico and against phytoplasmas of other groups; (2) defined reference lineages using ANI and a rooted phylogeny of single-copy orthologues; (3) tested whether the MLSA reproduces these lineages and assigns field isolates to them, in comparison with 16S rRNA classification; and (4) related the lineages to host range and geographic origin.

## Materials and methods

### Genome dataset and curation

Sixteen 16SrIII genome assemblies available in NCBI GenBank in October 2026 were downloaded with NCBI Datasets (Table 1), including the cassava oversprouting strains OVSP-S01 and OVSP-L01 (Oliveira et al., 2026). ‘*Ca.* P. phoenicium’ (16SrIX; GCA_056945795.1) and ‘*Ca.* P. oryzae’ HN2022 (16SrXI; GCA_027945135.1) were used as outgroups. Contigs with G+C content > 35% were screened by BLASTn against a local database of phytoplasma genomes and against NCBI nt; contigs without homology to phytoplasma genomes (≥ 50% query coverage), excluding those carrying rRNA genes (barrnap), were considered host-derived and removed. All assemblies were re-annotated with Prokka v1.15.6 (Seemann, 2014) using translation table 11. Completeness and contamination were estimated with CheckM v1.2.5 using the class *Mollicutes* marker set (Parks et al., 2015) and with BUSCO v6.0.0 (mollicutes_odb10) on predicted proteins.

### 16S rRNA analyses

Complete 16S rRNA genes (≥ 1,400 bp) were extracted with barrnap; one copy per genome was retained, plus additional copies only when they differed. Copies were screened for anomalies by locating the distribution of differences between paralogous copies and against other 16SrIII sequences; anomalous copies were replaced by the PCR-derived sequence of the same strain (ChTDIII, AY081817) or excluded. Virtual RFLP of the R16F2n/R16R2 region (Gundersen and Lee, 1996) with the 17 *i*PhyClassifier enzymes and similarity coefficients (fragments < 50 bp excluded) were computed with a custom script and confirmed with *i*PhyClassifier (Zhao et al., 2009). Full-length sequences were aligned with MAFFT L-INS-i (Katoh and Standley, 2013), pairwise identities were calculated excluding gapped positions, and a maximum-likelihood (ML) tree was inferred with IQ-TREE 3 (Wong et al., 2026).

### Average nucleotide identity and synteny

Pairwise ANI and aligned fractions were computed with skani (Shaw and Yu, 2023). Complete genomes representing three lineages (PR2021, LWB and OVSP-S01) were rotated to start at *dnaA*, aligned with MUMmer4 and visualised with pyGenomeViz. References for software not cited in the main text are provided in the Supplementary Material.

### Orthology and phylogenomics

Orthogroups were inferred with OrthoFinder (Emms and Kelly, 2019). Single-copy orthologues present in all 18 genomes were aligned at the protein level with MAFFT L-INS-i, back-translated to codon alignments with PAL2NAL (Suyama et al., 2006) and trimmed with trimAl (-automated1; Capella-Gutiérrez et al., 2009). Concatenated protein and codon supermatrices were analysed with IQ-TREE 3 using partitioned models selected by ModelFinder with partition merging (MFP+MERGE, -rcluster 10) and 1,000 ultrafast bootstrap (UFBoot) and SH-aLRT replicates. Alternative root positions were evaluated by constrained searches and the approximately unbiased (AU) test (Shimodaira, 2002). Gene and site concordance factors (gCF, sCF) were computed in IQ-TREE (Minh et al., 2020). Recombination was assessed with the PHI test (1,000 permutations; Bruen et al., 2006), and the SCG tree was re-inferred after excluding orthologues with significant signal.

### Marker selection and primer design

Candidate MLSA loci were selected among single-copy orthologues according to length, proportion of variable and parsimony-informative sites, pairwise identity and conserved flanking regions suitable for primer design. Of 36 candidate genes evaluated, primers were designed for seven: the five most informative (*cimH*, *mutM*, *ctpE*, *secY* and *gpmI*) and two conserved loci widely used in phytoplasma classification (*tuf* and *secA*), which allow comparison with published data. *ctpE* and *gpmI* were discarded because they did not amplify consistently from field isolates, and five loci were retained: *tuf* (elongation factor Tu), *secA* and *secY* (preprotein translocase subunits), *mutM* (formamidopyrimidine-DNA glycosylase) and *cimH* (citrate/malate transporter). Primers (Table 2) were evaluated with primer3-py for melting temperature, hairpins and dimers, and *in silico* against all genomes by locating the best binding site and counting total and 3′-terminal (last five bases) mismatches.

### Plant material, PCR and sequencing

Five isolates previously characterised by 16S rRNA analysis were analysed (Table 1): BidPhy (16SrIII-X, *Bidens subalternans*), FodderBeet (16SrIII-J, *Beta vulgaris* var. *rapacea*), CassDWB (16SrIII-J, *Manihot esculenta*; Fernández et al., 2018), PYWB (16SrIII-B, *Prunus domestica*; Bongiorno et al., 2020) and CaesLL (16SrIII-B, *Erythrostemon gilliesii*, syn. *Caesalpinia gilliesii*; Alessio et al., 2025). Two further isolates (ArPY and HetPhy) failed to amplify, consistent with low phytoplasma titre in stored DNA. Total DNA was extracted as described previously (Fernández et al., 2018). Each locus was amplified in 25-µL reactions containing 5 µL of 5× Q5 reaction buffer, 2.5 µL of 10 mM dNTPs, 2.5 µL of each primer (10 µM), 0.25 µL of Q5 High-Fidelity DNA polymerase (New England Biolabs), 0.75 µL of template DNA and nuclease-free water. Cycling conditions were 98 °C for 30 s; 30 cycles of 98 °C for 10 s, 60–63 °C for 30 s depending on the locus (Table 2) and 72 °C for 30 s; and a final extension at 72 °C for 2 min. Products were visualised in 1% agarose gels and purified with AMPure XP magnetic beads (Beckman Coulter) before sequencing. Specificity was evaluated with DNA from seven 16SrIII isolates (including BellVir, 16SrIII-J, maintained in periwinkle) and from phytoplasmas of other groups occurring in the same regions: 16SrVII-C (ArAWB, alfalfa), 16SrXIII-G (‘*Ca.* P. meliae’ ChTYXIII, chinaberry) and 16SrI (BidPhy-Cba, an isolate from *Bidens subalternans* distinct from the 16SrIII-X isolate BidPhy); the 16S rRNA gene was amplified as a positive control. Amplicons were sequenced in both directions (Sanger; Macrogen, Seoul, South Korea). Consensus sequences were assembled and inspected manually; apparent indels in homopolymers were verified against chromatograms using peak spacing and reading frame.

### MLSA

For each locus, orthologues were extracted from genome assemblies by BLASTn with the isolate consensus sequences (tBLASTn for outgroups), aligned with MAFFT and trimmed to the region covered by all Sanger sequences. Because outgroup branches were very long (> 0.7 substitutions/site) and caused long-branch attraction with short loci, outgroups were excluded from MLSA analyses, and MLSA trees were rooted at the position inferred from the SCG phylogeny. ML trees were inferred for each locus and for the concatenated alignment (partitioned, MFP+MERGE). Congruence with the SCG trees was assessed with normalised Robinson–Foulds distances (nRF) and with the SH and AU tests on the genomes shared by both datasets, after collapsing near-zero-length branches. The five MLSA loci represent ∼1% of the SCG alignment, so their contribution to the SCG topology is negligible.

## Results

### Genome dataset, curation and quality

The 16 assemblies ranged from complete genomes (PR2021, LWB, OVSP-S01, OVSP-L01; 1–2 replicons) to drafts of up to 272 contigs (Table 1). Thirteen ChTDIII contigs (57.3 kb; G+C 35–44%) lacked homology to phytoplasma genomes or NCBI nt and were removed, reducing this assembly from 790.5 to 733.2 kb. After curation, genome sizes were 583.8–758.2 kb, G+C content 26.7–28.7%, and 513–805 CDS were predicted. CheckM completeness reached 97.1% for complete genomes, the maximum attainable with this marker set; 12 assemblies were within 2% of this value, MA1 and Vc33 showed 91.4% and 86.4%, and contamination was ≤ 0.95%.

### 16S rRNA classification and anomalous copies

Virtual RFLP confirmed published subgroups and assigned previously unclassified genomes (Table 1): CanS4 shared the 16SrIII-F pattern, OVSP-S01 and OVSP-L01 the 16SrIII-B pattern, and JR1 and PR2021 a pattern most similar to 16SrIII-A (F = 0.99), although JR1 had been reported as 16SrIII-H. The five field isolates showed patterns identical to their published subgroups. Two genome-derived 16S rRNA copies were anomalous. In ChTDIII, positions ∼60–440 (V2–V3) diverged from all 16SrIII sequences (88% identity), whereas the remainder was identical to 16SrIII-B; the PCR-derived sequence (AY081817) showed the 16SrIII-B pattern (F = 1.00) and was used instead. In OVSP-L01, the rrn2 copy contained a 105-bp block (positions 285–389, V2–V3; 62 differences including a 10-nt insertion), while the remaining ∼1,415 bp were identical to rrn1; this copy, which had produced a spurious RFLP pattern (F = 0.90), was excluded. Excluding it, full-length 16S rRNA identity among all 16SrIII genomes was 98.89–99.87%, above the 98.65% species threshold, and the 16S tree was poorly resolved, with most nodes below 80% support (Fig. S1).

### ANI delineates four lineages

ANI values (Fig. 2) defined three clusters at ≥ 97% ANI and an isolated strain: Lineage I (CX, 6A1, ZT3-1, JR1, PR2021, PDA15; 97.4–99.9%), Lineage II (CanS4, MW1, VAC, MA1; 97.0–99.9%), Lineage III (ChTDIII, CicWB-2022, Vc33, OVSP-S01, OVSP-L01; 95.1–99.6%) and Lineage IV (LWB, 16SrIII-X; ≤ 95.6% to any genome). Lineage III shared 90.8–93.9% ANI with Lineages I and II. Aligned fractions between lineages were 52–71%.

### Rooted SCG phylogeny

OrthoFinder identified 223 single-copy orthologues shared by the 16 ingroup and two outgroup genomes. Protein and codon trees recovered the four lineages with SH-aLRT/UFBoot 100/100 (Fig. 2, Fig. S2). The root was placed on the branch leading to Lineage III (100/100 in both trees), and LWB was sister to Lineages I + II. Alternative root positions, with the outgroups constrained as sister to LWB, to Lineage II or to Lineage I, were significantly rejected (AU test, p ≤ 2.4 × 10^−4^; ΔlnL 191.6–245.8). Gene and site concordance factors supported the three multi-strain lineages (gCF 65.6–73.2%; sCF 75.7–88.6%), whereas the root position and the LWB–Lineage I + II relationship, although fully supported by concatenation and the AU test, showed substantial gene-tree discordance (gCF 29.5–31.7%), as expected for short internal branches. Within Lineage III, relationships among 16SrIII-J, OVSP and ChTDIII differed between protein and codon trees and were unresolved (gCF 14.3%; sCF 36.2% versus 35.0% and 28.8% for the alternatives). Twenty-one of 224 orthologues (9.4%) showed recombination signal (PHI p < 0.05), close to the number expected by chance; a tree inferred without them (203 loci; 195,312 nt) was identical to the full tree (RF = 0).

### Genomic context of MLSA loci and primer evaluation

In the three complete genomes, the five loci were single-copy and located in high-identity regions (Fig. 1). The chromosome ends around *dnaA* were collinear across lineages, whereas the central region showed rearrangements and inversions; *secA* and *cimH* retained the same relative position, while the blocks carrying *tuf*, *secY* and *mutM* were rearranged between lineages. In silico, the five loci were single-copy and fully covered in all 16 genomes; primers matched 11 genomes without mismatches, and the remaining mismatches (1–2 in LWB, OVSP-S01/L01 and ZT3-1) were outside the 3′ end except for one in *mutM*-R of ZT3-1 (Table 2).

**Fig. 1.**
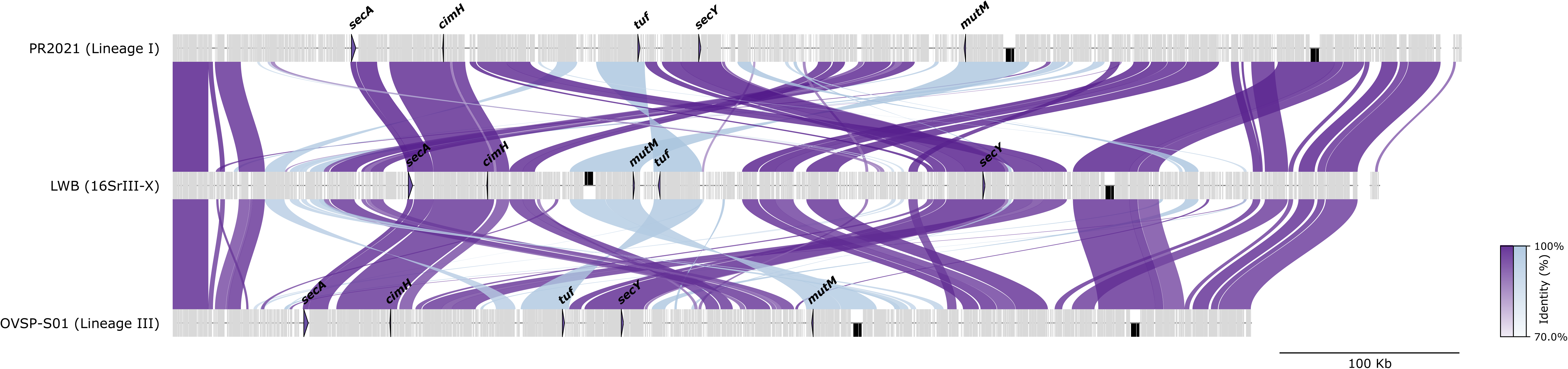
Genome synteny among complete genomes of three X-disease lineages (PR2021, Lineage I; LWB, 16SrIII-X; OVSP-S01, Lineage III). Chromosomes were rotated to start at *dnaA*. Links show MUMmer alignments ≥ 1 kb coloured by nucleotide identity (dark, same strand; light, inverted). The five MLSA loci are highlighted; black boxes, rRNA operons.

**Fig. 2.**
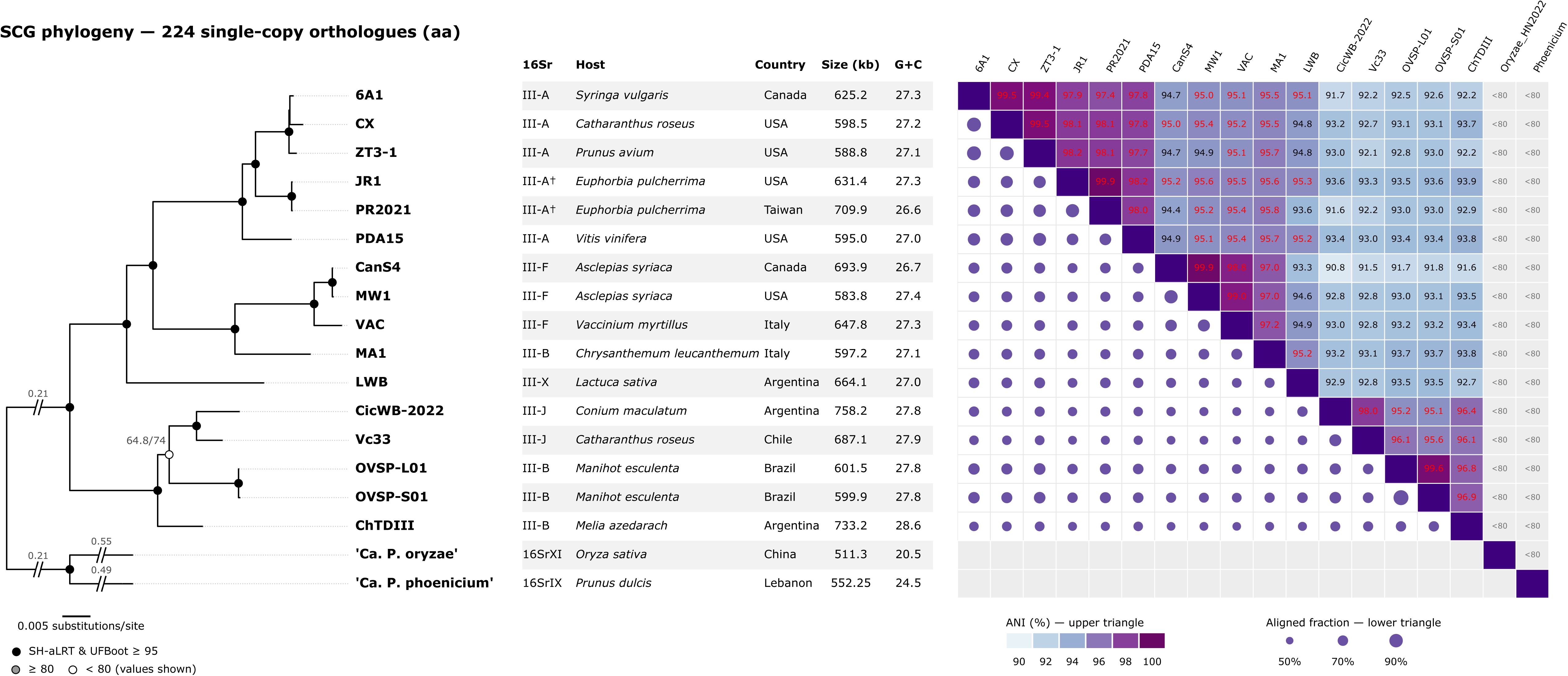
Maximum-likelihood phylogeny of 16 X-disease group genomes based on 223 single-copy orthologues (protein alignment), rooted with ‘*Ca.* P. phoenicium’ and ‘*Ca.* P. oryzae’ (outgroup branches truncated; real length shown), with strain metadata and pairwise ANI (upper triangle) and aligned fraction (lower triangle, circle size). Node circles: filled, SH-aLRT and UFBoot ≥ 95; grey, ≥ 80; open, < 80 (values shown). Cells framed: ANI ≥ 95%.

### Specificity of MLSA primers

All loci were amplified from the seven 16SrIII isolates tested, representing subgroups 16SrIII-B, -J and -X (Table 5). No amplification of *tuf*, *secA*, *secY* or *mutM* was obtained from phytoplasmas of groups 16SrI, 16SrVII and 16SrXIII, whereas *cimH* produced an amplicon of the expected size from the 16SrI isolate BidPhy-Cba. The 16S rRNA gene was amplified from all samples.

### MLSA

Amplicons were obtained from the five sequenced isolates for all loci. Apparent compensating indels in *mutM* of FodderBeet and PYWB were base-calling artefacts in homopolymers (split peaks at half the median peak spacing, or low-quality read ends) and were corrected, restoring the reading frame. Loci differed in informativeness (Table 3), with *cimH* and *mutM* the most variable and *tuf* and *secA* the most conserved; none showed recombination signal (PHI p = 0.14–0.99). The concatenated alignment (21 sequences; 2,870 nt; 358 parsimony-informative sites) recovered the same lineages as the SCG phylogeny (Fig. 3). nRF distances to the SCG trees were 0.077 (protein) and 0.154 (codon) for the concatenation and 0.08–0.38 for single loci. The SCG protein topology was statistically indistinguishable from the MLSA tree (ΔlnL = 0.03; SH test p = 0.51), whereas the codon topology, which differs in the position of ChTDIII relative to OVSP, was marginally rejected (ΔlnL = 12.9; p-AU = 0.029). Deep nodes were supported by four or five loci, whereas nodes within Lineage III depended on one or two loci (Fig. 3, concordance panel). BidPhy was identical to LWB at all loci, FodderBeet was identical to CicWB-2022, CassDWB grouped within 16SrIII-J, and PYWB grouped with ChTDIII (76.8/73). CaesLL, a 16SrIII-B isolate, grouped within the 16SrIII-J clade (96–97/96) as sister to CassDWB (80.4/79). Thus, 16SrIII-B strains occupied three positions within Lineage III and one in Lineage II.

**Fig. 3.**
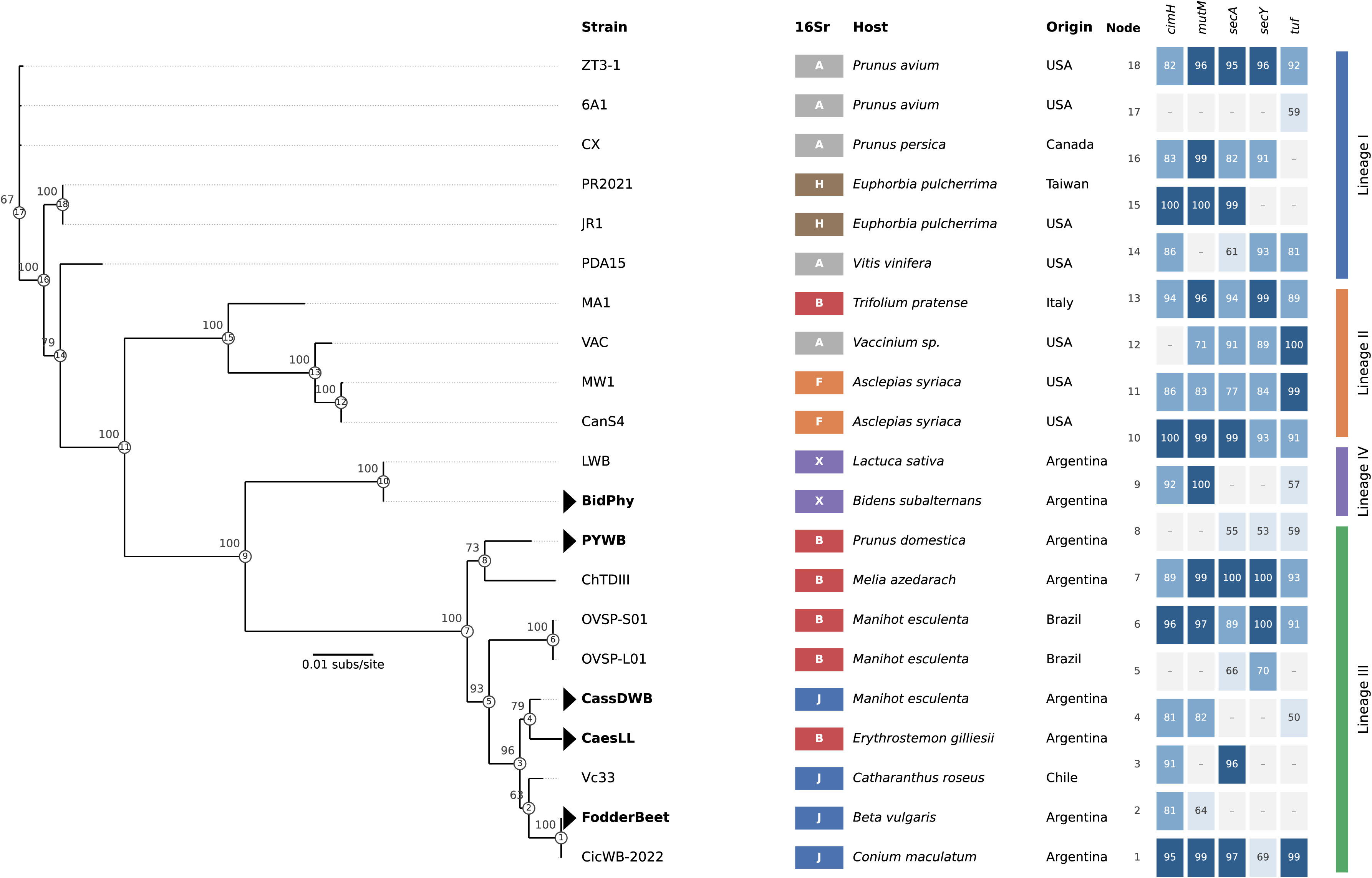
Concatenated MLSA tree (*tuf*, *secA*, *secY*, *mutM*, *cimH*; 2,870 nt) including five isolates sequenced in this study (triangles), rooted at the position inferred from the SCG phylogeny, with gene-concordance matrix showing the UFBoot support of each node in single-locus trees.

### Host range and geographic origin of lineages

The lineages showed contrasting host and geographic ranges (Table 4). Lineage III, represented exclusively by South American strains, infected hosts from six plant families, including staple (cassava), fruit (plum) and forage (fodder beet) crops, an ornamental and forest tree (chinaberry), a native shrub and a weed. Cassava hosted 16SrIII-B strains in Brazil and a 16SrIII-J strain in Argentina, all belonging to Lineage III, whereas *Prunus* hosted Lineage I in North America (sweet cherry) and Lineage III in Argentina (plum). Lineage I comprised strains from North America and Asia infecting four plant families, including fruit crops (cherry, grapevine) and ornamentals; Lineage II included strains from North America and Europe, mostly from wild plants; and Lineage IV was found in Asteraceae in Argentina.

## Discussion

We developed a genome-guided MLSA scheme for X-disease group phytoplasmas based on five single-copy housekeeping loci selected from genome-wide orthology. The loci are present in all available genomes, lie in conserved regions of the chromosome, show no evidence of recombination and can be amplified directly from field samples by conventional PCR. The concatenated analysis was statistically compatible with the phylogeny inferred from 223 single-copy orthologues, so that sequencing ∼2.9 kb is sufficient to assign a strain to the genomic lineages defined by whole-genome data. This validation against genome-scale relationships distinguishes the scheme from marker sets chosen *a priori*, and extends to the X-disease group the genome-guided approach proposed for 16SrI phytoplasmas (Cho et al., 2020), adding a rooted reference phylogeny, statistical tests of congruence and amplification from field samples.

The scheme resolves diversity that the 16S rRNA gene does not detect. Full-length 16S rRNA sequences were 98.89–99.87% identical across the group and produced a poorly resolved tree, whereas the five loci separated four lineages with full support. Subgroup assignment by RFLP was not a reliable proxy for these lineages: strains with identical 16SrIII-B patterns occurred in two lineages and, within the South American lineage, in three different positions, including an isolate nested among 16SrIII-J strains. These results generalise the divergence of MA1 from Argentinian 16SrIII-B strains previously observed with *imp* and *idpA* (Alessio et al., 2025). CaesLL grouped with ChTDIII in those genes but with 16SrIII-J in our housekeeping loci; because *imp* is under positive selection, markers of that type are informative about host or vector interactions but less suitable for lineage assignment.

For diagnosis and epidemiology, the lineage rather than the 16Sr subgroup appears to be the relevant unit. Strains assigned to 16SrIII-B and 16SrIII-J in South America share a single genomic background with a broad host range that spans crops, trees, native plants and weeds, consistent with weeds and native plants acting as reservoirs for crop infections. Conversely, the same host genus can harbour unrelated lineages, as shown for *Prunus*. This interpretation is supported by an independent multilocus analysis of 16SrIII genomes based on *secY*, *secA*, *nusA*, *tuf* and ribosomal protein genes, which recovered groupings consistent with the lineages defined here and likewise placed the 16SrIII-B strains MA1 and ChTDIII in different clades (Pellegrinetti et al., 2026). Lineage-level variation is also relevant in North America, where ‘*Ca.* P. pruni’ lineages differ in regional distribution and in the symptoms they induce on cherry (Molnar et al., 2024), and geographic structure within subgroups has been reported between North and South American strains (Pérez-López et al., 2017). The scheme used by Molnar et al. (2024) shares *secY*, *secA* and *tuf* with ours but adds the hypervariable *imp* gene to resolve strains within ‘*Ca.* P. pruni’ *sensu stricto*; combining both approaches would allow typing from lineages down to strains. Lineage-level typing could therefore improve surveys, the identification of reservoirs and vectors, and risk assessment for crops such as cassava and stone fruits, where 16S-based assignment alone does not discriminate between unrelated lineages.

The lineages recognised here also bear on classification. The South American lineage shares < 95% ANI with ‘*Ca.* P. pruni’ *sensu stricto*, although 16S rRNA identity exceeds the 98.65% threshold (Bertaccini et al., 2022), a discordance between 16S-based and genome-based delimitation also reported for other phytoplasma groups (Cho et al., 2020; Kirdat et al., 2023). 16SrIII-J strains of this lineage were previously proposed as a distinct taxon (Fernández et al., 2019), and our results extend the lineage to South American 16SrIII-B strains. Lineage IV (16SrIII-X) also shares ≤ 95.6% ANI with all other genomes, over aligned fractions of only 52–71%. We refrain from formal proposals here, as the taxonomic status of these lineages is being addressed in a separate genome-based study.

## Data availability

Genome accessions are listed in Table 1. Sequences of the five MLSA loci from the five field isolates were deposited in GenBank under accessions QB286830-QB286854 (Table S3). Alignments, phylogenetic trees and analysis outputs files are available at Zenodo (https://doi.org/10.5281/zenodo.24204081).

## Acknowledgements

This research was funded by the National Institute of Agricultural Technology (INTA) through Projects 2023-PD-L03-I084 and 2023-PD-L01-I085, and by Fundación ArgenINTA.

## Author contributions

F.D.F.: conceptualisation, methodology, formal analysis, bioinformatics, funding acquisition, writing – original draft. V.A.B.: investigation (PCR and Sanger sequencing), writing – review and editing. L.R.C.: supervision, writing – review and editing. All authors read and approved the final manuscript.

## Supporting Information legends

**Table S1.** Recombination test (PHI, 1,000 permutations) for the 224 single-copy orthologues used in the SCG phylogeny.

**Table S2.** In silico evaluation of MLSA primers against 16 genome assemblies (total mismatches and mismatches within the five 3′-terminal bases).

**Table S3.** GenBank accession numbers of the MLSA loci sequenced in this study (QB286830–QB286854).

**Figure S1.**
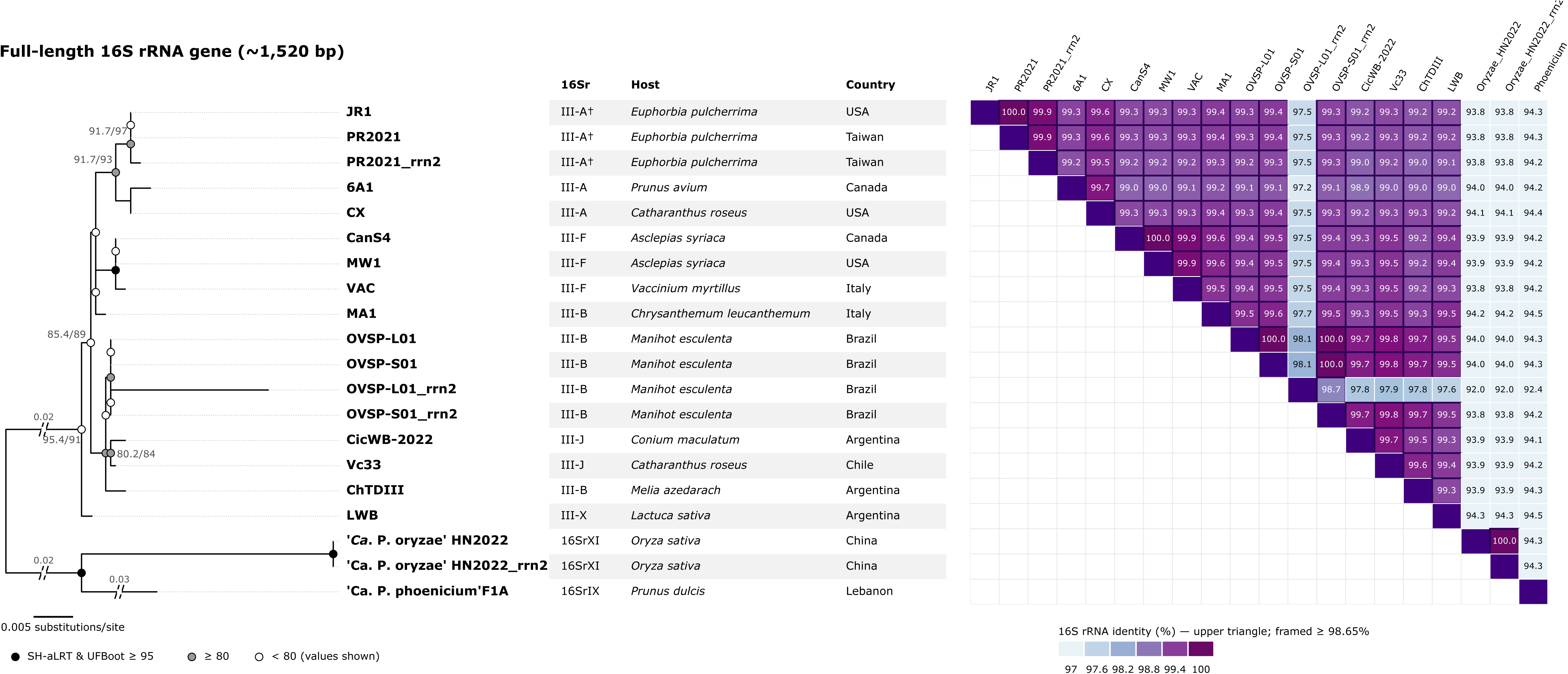
Maximum-likelihood tree and pairwise identity of full-length 16S rRNA genes; cells framed ≥ 98.65% (Bertaccini et al., 2022).

**Figure S2.**
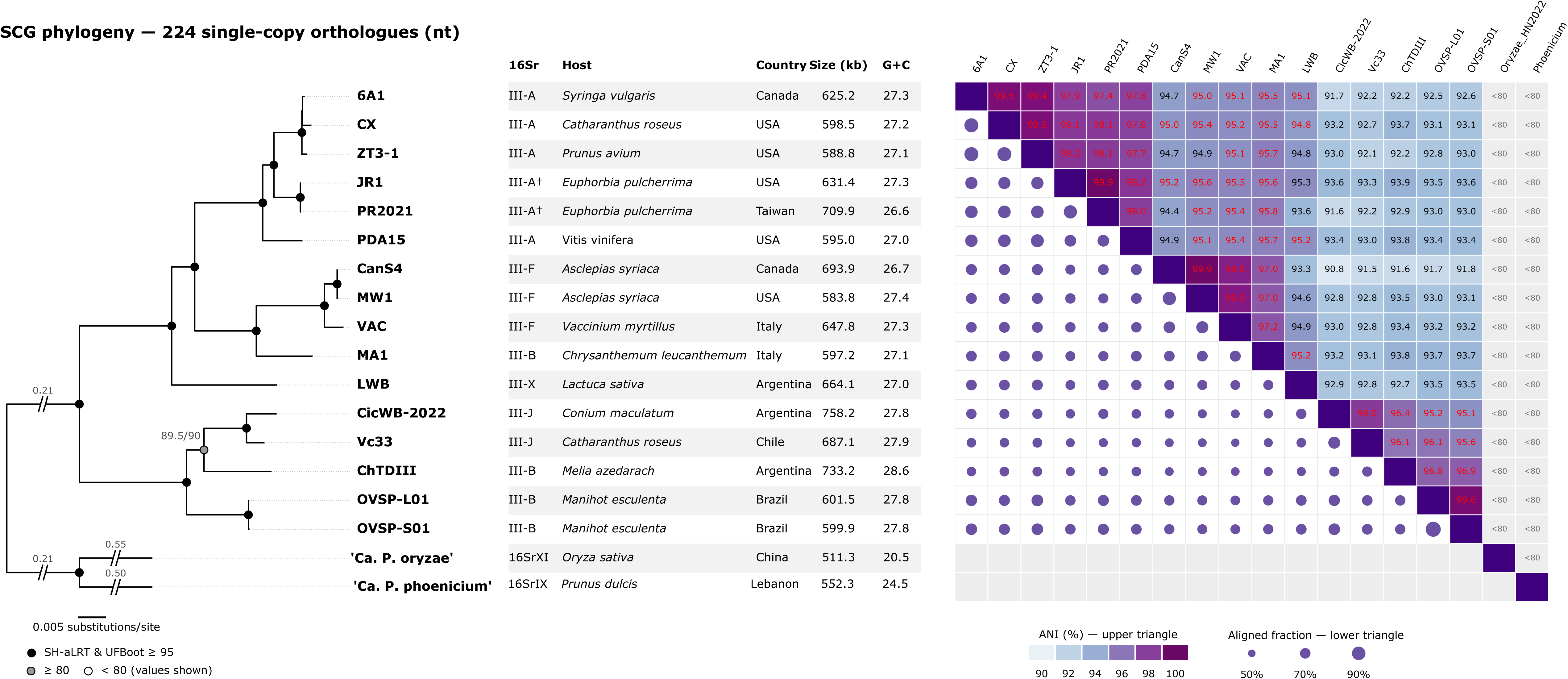
Maximum-likelihood SCG phylogeny based on the codon alignment of 223 single-copy orthologues, rooted with ‘*Ca.* P. phoenicium’ and ‘*Ca.* P. oryzae’, with pairwise ANI.

**Figure S3.**
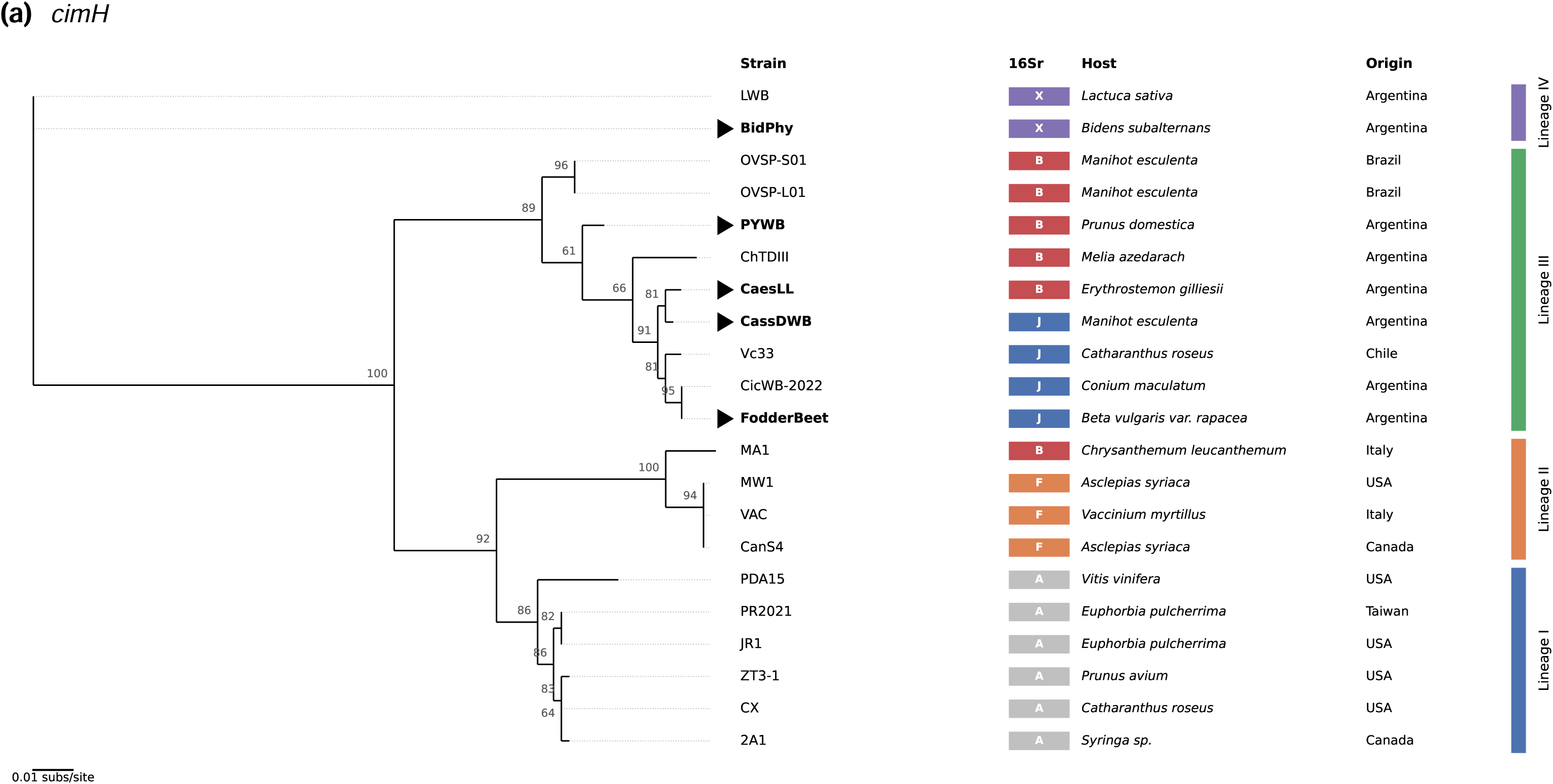
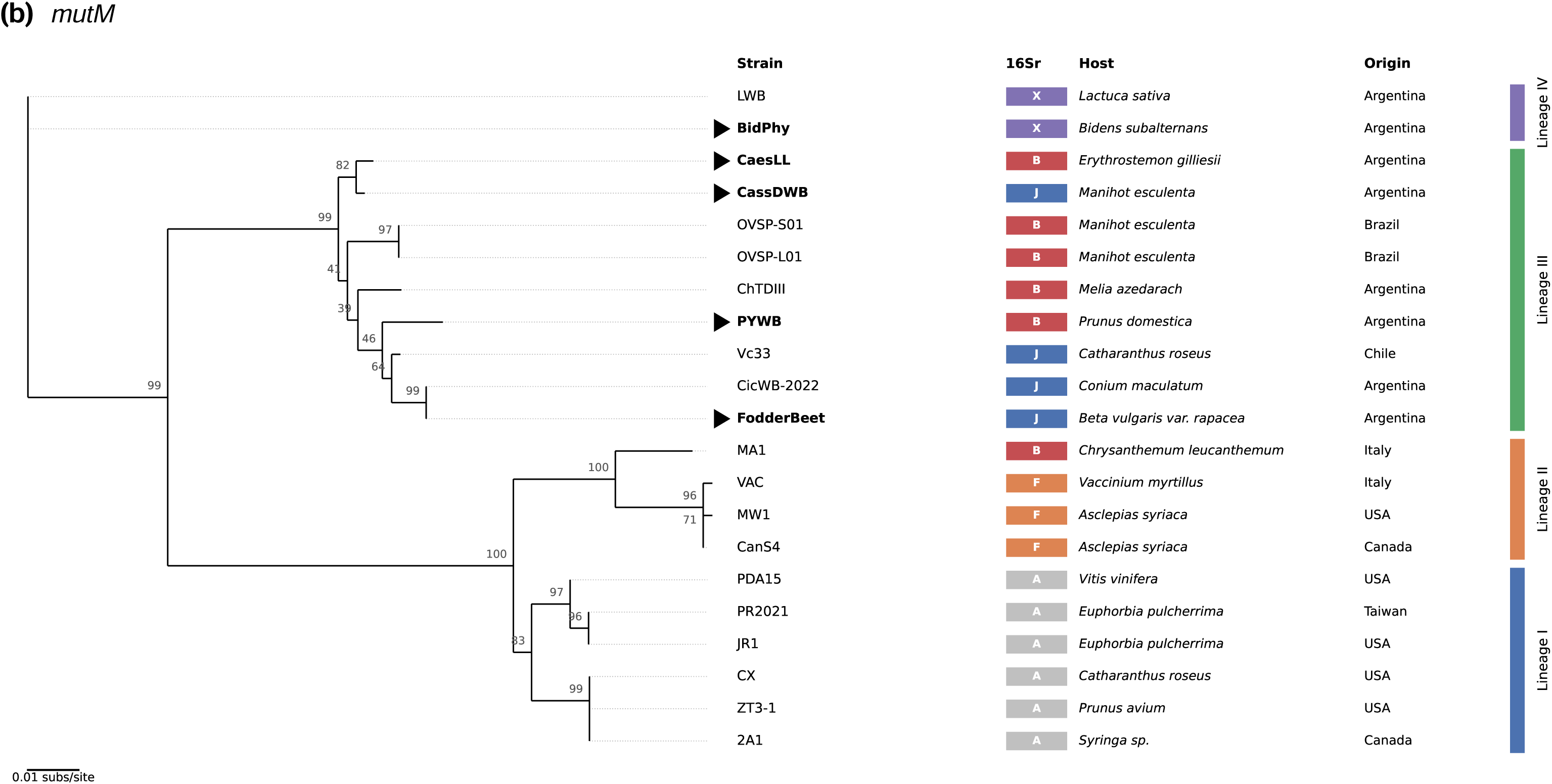
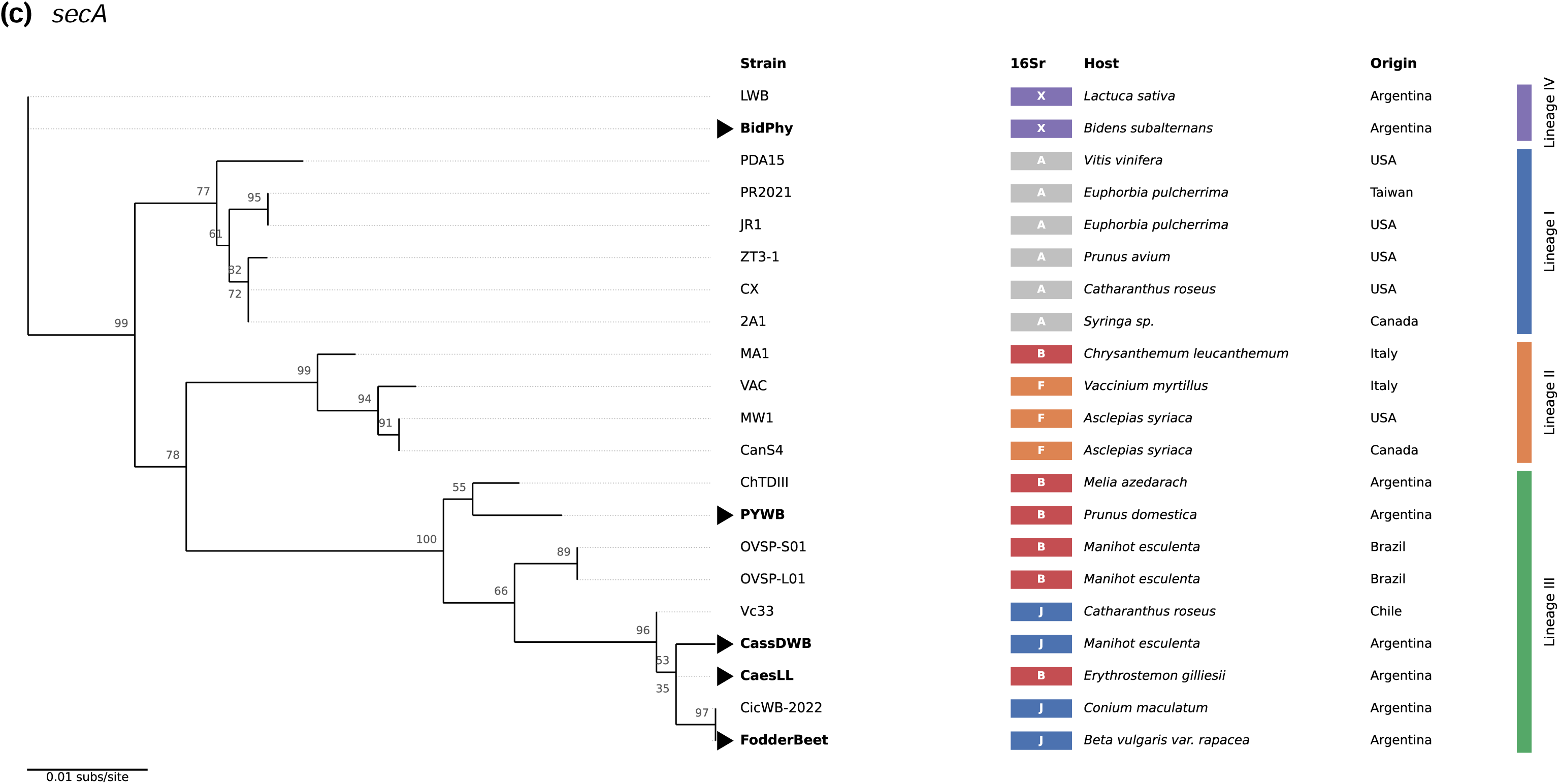
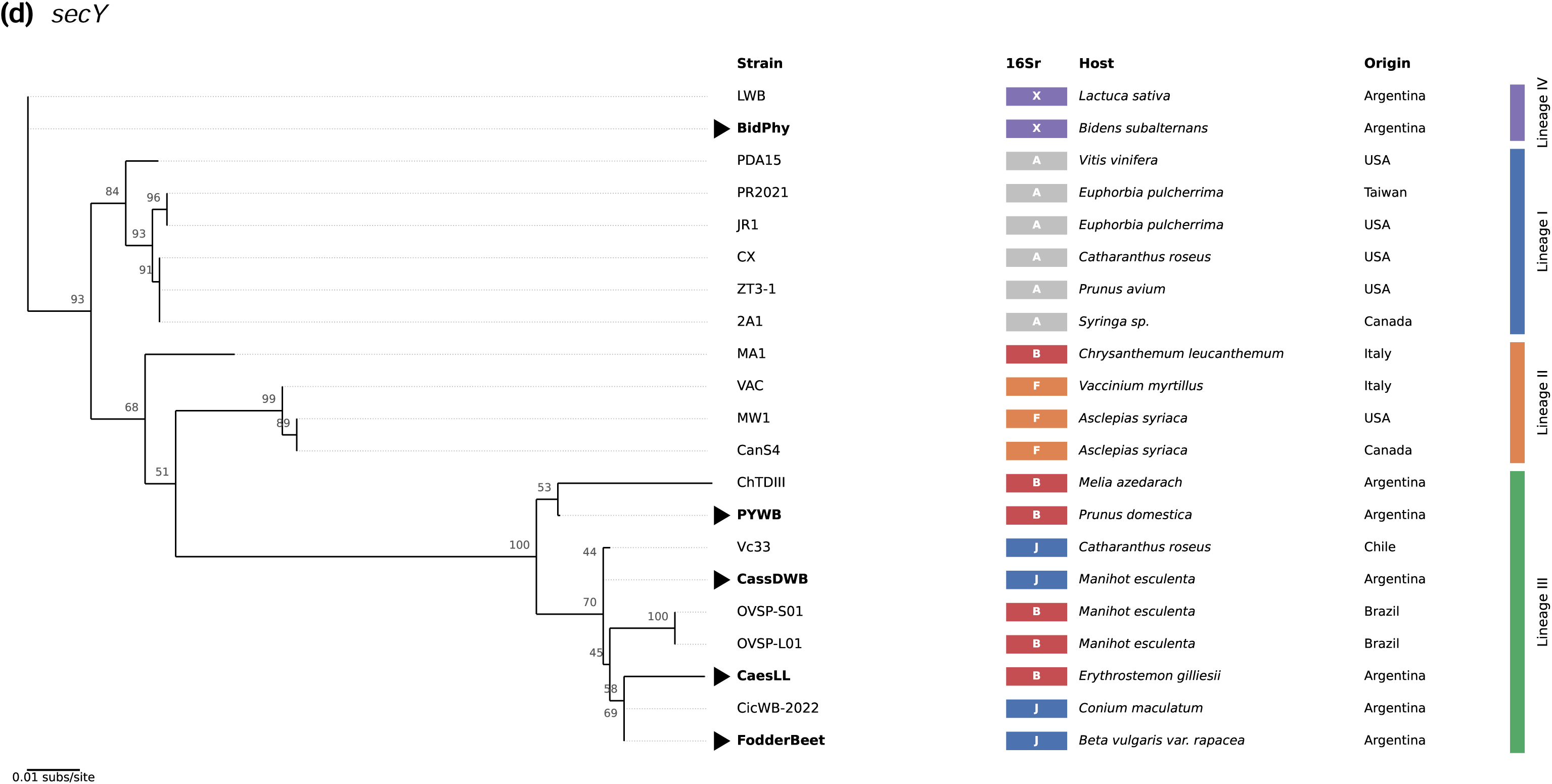
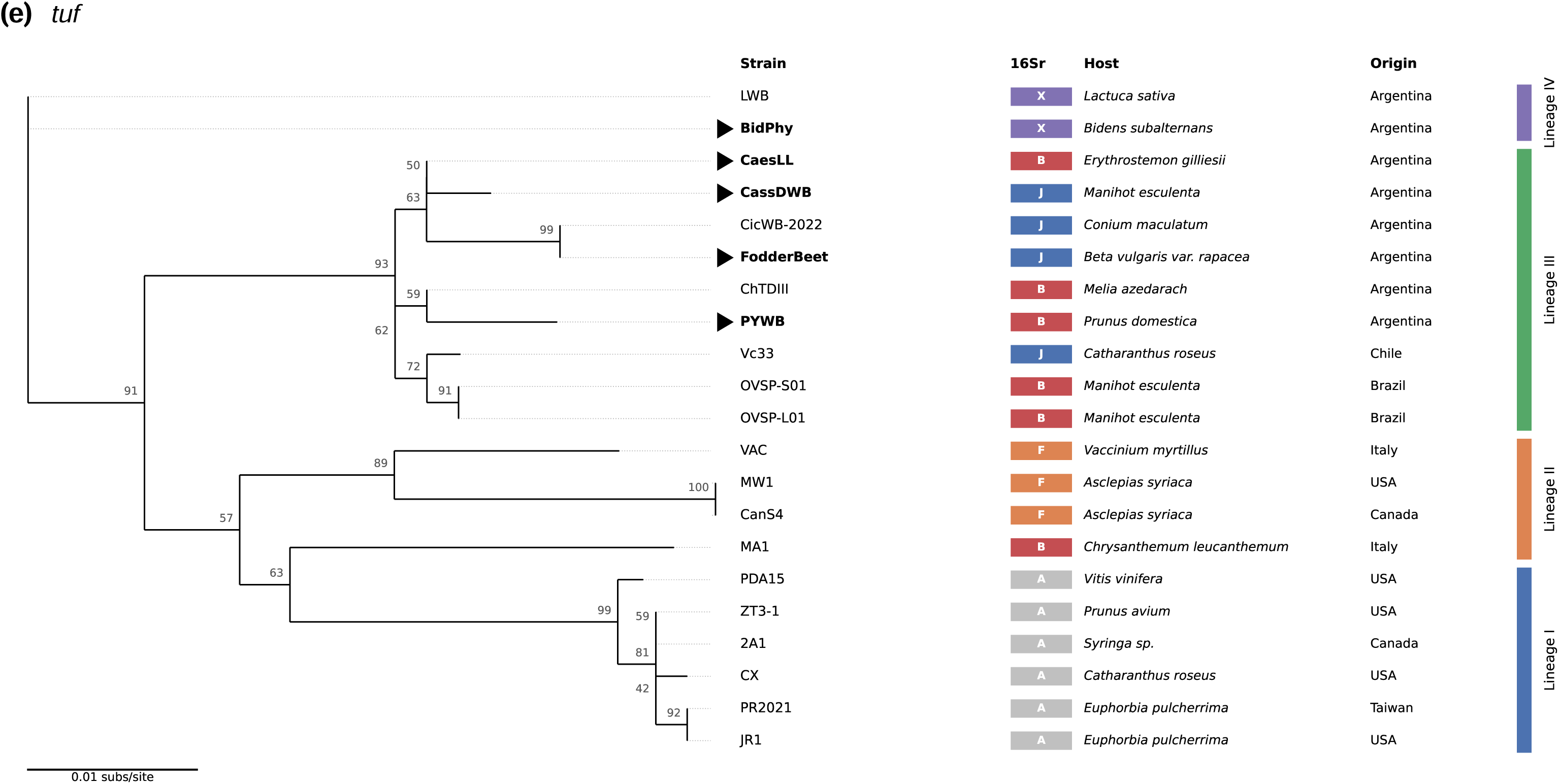
Single-locus maximum-likelihood trees for (a) *cimH*, (b) *mutM*, (c) *secA*, (d) *secY* and (e) *tuf* (21 sequences each), rooted at the position inferred from the SCG phylogeny. Node labels are UFBoot values; triangles and bold type mark the field isolates sequenced in this study. Columns give the 16Sr subgroup, host and country of each strain; bars on the right delimit lineages I–IV. The four lineages are recovered as contiguous groups by every locus, although support for internal nodes within lineage III varies among loci. Software references not cited in the main text are listed in the Supporting Information.

