## Supplementary material for "A genome-guided multilocus sequence analysis scheme to detect and resolve diversity within the X-disease phytoplasma group (16SrIII)": Supp_info

**Table S1. Recombination test (PHI, 1,000 permutations) for the 224 single-copy orthologues used in the SCG phylogeny.**

Provided as a separate Excel file (TableS1_PHI.xlsx). Twenty-one of the 224 orthologues showed significant recombination signal (p < 0.05); excluding them did not change the SCG topology (RF = 0).

**Table S2. In silico evaluation of MLSA primers against 16 genome assemblies: total mismatches (mismatches within the five 3′-terminal bases).**

| **Genome** | ***tuf* F** | ***tuf* R** | ***secA* F** | ***secA* R** | ***secY* F** | ***secY* R** | ***mutM* F** | ***mutM* R** | ***cimH* F** | ***cimH* R** |
| --- | --- | --- | --- | --- | --- | --- | --- | --- | --- | --- |
| 2A1 | 0 (0) | 0 (0) | 0 (0) | 0 (0) | 0 (0) | 0 (0) | 0 (0) | 0 (0) | 0 (0) | 0 (0) |
| CX | 0 (0) | 0 (0) | 0 (0) | 0 (0) | 0 (0) | 0 (0) | 0 (0) | 0 (0) | 0 (0) | 0 (0) |
| ZT3-1 | 0 (0) | 0 (0) | 0 (0) | 0 (0) | 0 (0) | 0 (0) | 0 (0) | 1 (1) | 0 (0) | 0 (0) |
| PDA15 | 0 (0) | 0 (0) | 0 (0) | 0 (0) | 0 (0) | 0 (0) | 0 (0) | 0 (0) | 0 (0) | 0 (0) |
| JR1 | 0 (0) | 0 (0) | 0 (0) | 0 (0) | 0 (0) | 0 (0) | 0 (0) | 0 (0) | 0 (0) | 0 (0) |
| PR2021 | 0 (0) | 0 (0) | 0 (0) | 0 (0) | 0 (0) | 0 (0) | 0 (0) | 0 (0) | 0 (0) | 0 (0) |
| CanS4 | 0 (0) | 0 (0) | 0 (0) | 0 (0) | 0 (0) | 0 (0) | 0 (0) | 0 (0) | 0 (0) | 0 (0) |
| MW1 | 0 (0) | 0 (0) | 0 (0) | 0 (0) | 0 (0) | 0 (0) | 0 (0) | 0 (0) | 0 (0) | 0 (0) |
| VAC | 0 (0) | 0 (0) | 0 (0) | 0 (0) | 0 (0) | 0 (0) | 0 (0) | 0 (0) | 0 (0) | 0 (0) |
| MA1 | 0 (0) | 0 (0) | 0 (0) | 0 (0) | 0 (0) | 0 (0) | 0 (0) | 0 (0) | 0 (0) | 0 (0) |
| ChTDIII | 0 (0) | 0 (0) | 0 (0) | 0 (0) | 0 (0) | 0 (0) | 0 (0) | 0 (0) | 0 (0) | 0 (0) |
| OVSP-S01 | 0 (0) | 0 (0) | 0 (0) | 1 (0) | 0 (0) | 0 (0) | 1 (0) | 1 (0) | 0 (0) | 0 (0) |
| OVSP-L01 | 0 (0) | 0 (0) | 0 (0) | 1 (0) | 0 (0) | 0 (0) | 1 (0) | 1 (0) | 0 (0) | 0 (0) |
| CicWB-2022 | 0 (0) | 0 (0) | 0 (0) | 0 (0) | 0 (0) | 0 (0) | 0 (0) | 0 (0) | 0 (0) | 0 (0) |
| Vc33 | 0 (0) | 0 (0) | 0 (0) | 0 (0) | 0 (0) | 0 (0) | 0 (0) | 0 (0) | 0 (0) | 0 (0) |
| LWB | 0 (0) | 0 (0) | 0 (0) | 0 (0) | 1 (0) | 0 (0) | 0 (0) | 0 (0) | 2 (0) | 2 (0) |

All five loci were single-copy with full-length coverage in all genomes. Degenerate positions were evaluated as IUPAC codes.

**Table S3. GenBank accession numbers of the MLSA loci sequenced in this study.**

| **Isolate** | **16Sr** | ***tuf*** | ***secA*** | ***secY*** | ***mutM*** | ***cimH*** |
| --- | --- | --- | --- | --- | --- | --- |
| BidPhy | III-X | QB286830 | QB286834 | QB286838 | QB286842 | QB286844 |
| FodderBeet | III-J | QB286831 | QB286835 | QB286839 | QB286848 | QB286845 |
| CassDWB | III-J | QB286832 | QB286836 | QB286840 | QB286843 | QB286846 |
| PYWB | III-B | QB286833 | QB286837 | QB286841 | QB286849 | QB286847 |
| CaesLL | III-B | QB286850 | QB286851 | QB286852 | QB286853 | QB286854 |

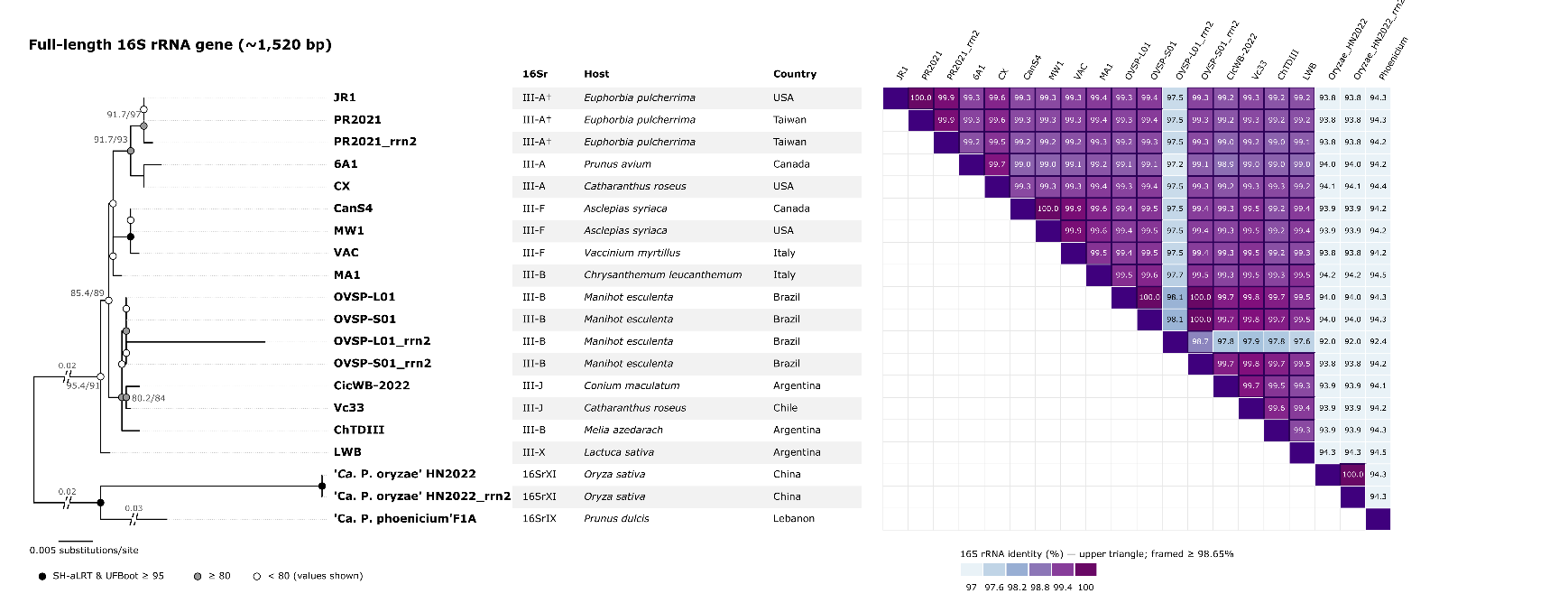

**Fig. S1.** Maximum-likelihood tree and pairwise identity of full-length 16S rRNA genes (~1,520 bp) of 16SrIII genomes and outgroups; cells framed ≥ 98.65% (Bertaccini et al., 2022). Anomalous copies (ChTDIII assembly-derived 16S; OVSP-L01 rrn2) were replaced or excluded.

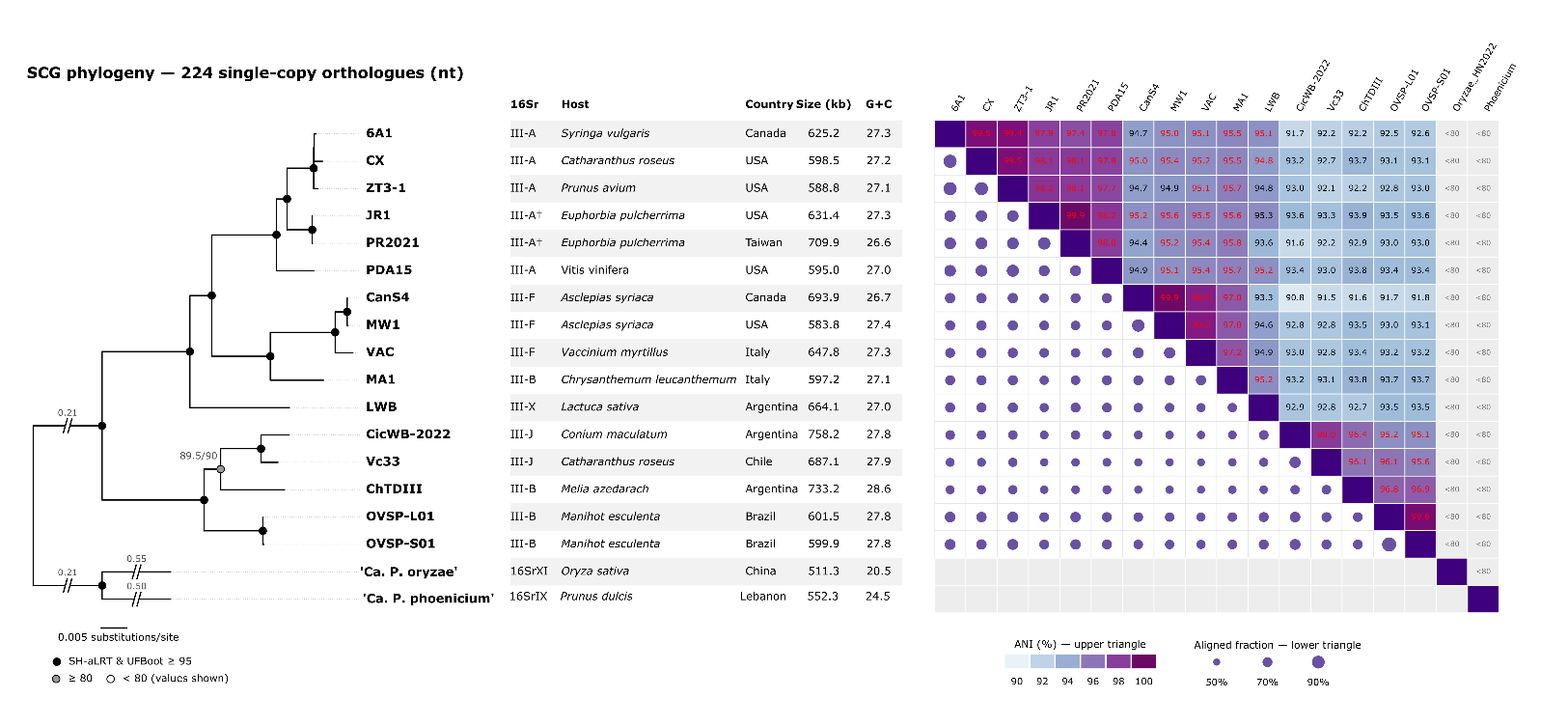

**Fig. S2.** Maximum-likelihood SCG phylogeny based on the codon alignment of 224 single-copy orthologues, rooted with ‘*Ca.* P. phoenicium’ and ‘*Ca.* P. oryzae’, with pairwise ANI (values ≥ 95 in red)

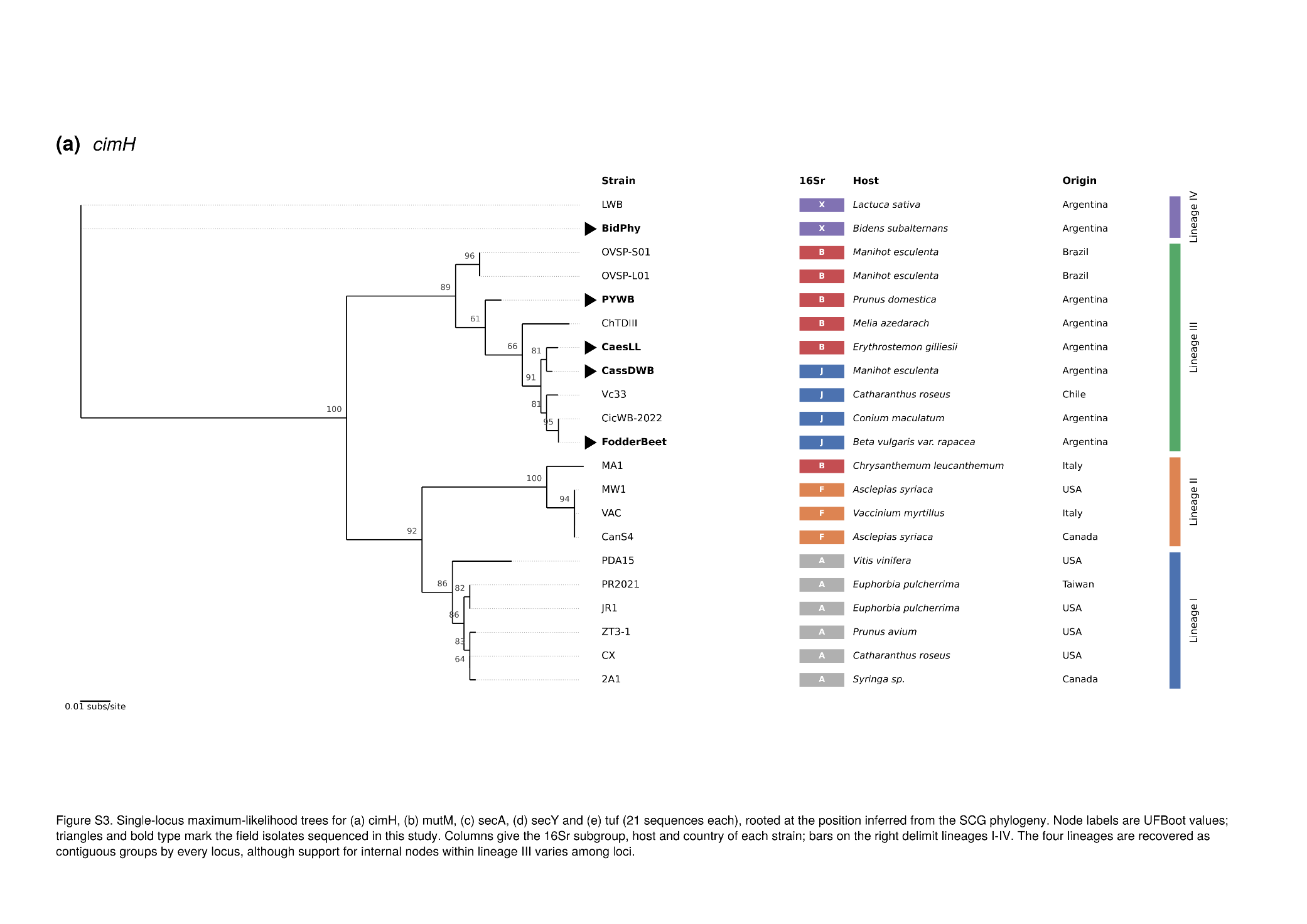

**Figure S3.** Single-locus maximum-likelihood trees for (a) *cimH*, (b) *mutM*, (c) *secA*, (d) *secY* and (e) *tuf* (21 sequences each), rooted at the position inferred from the SCG phylogeny. Node labels are UFBoot values; triangles and bold type mark the field isolates sequenced in this study. Columns give the 16Sr subgroup, host and country of each strain; bars on the right delimit lineages I–IV. The four lineages are recovered as contiguous groups by every locus, although support for internal nodes within lineage III varies among loci.

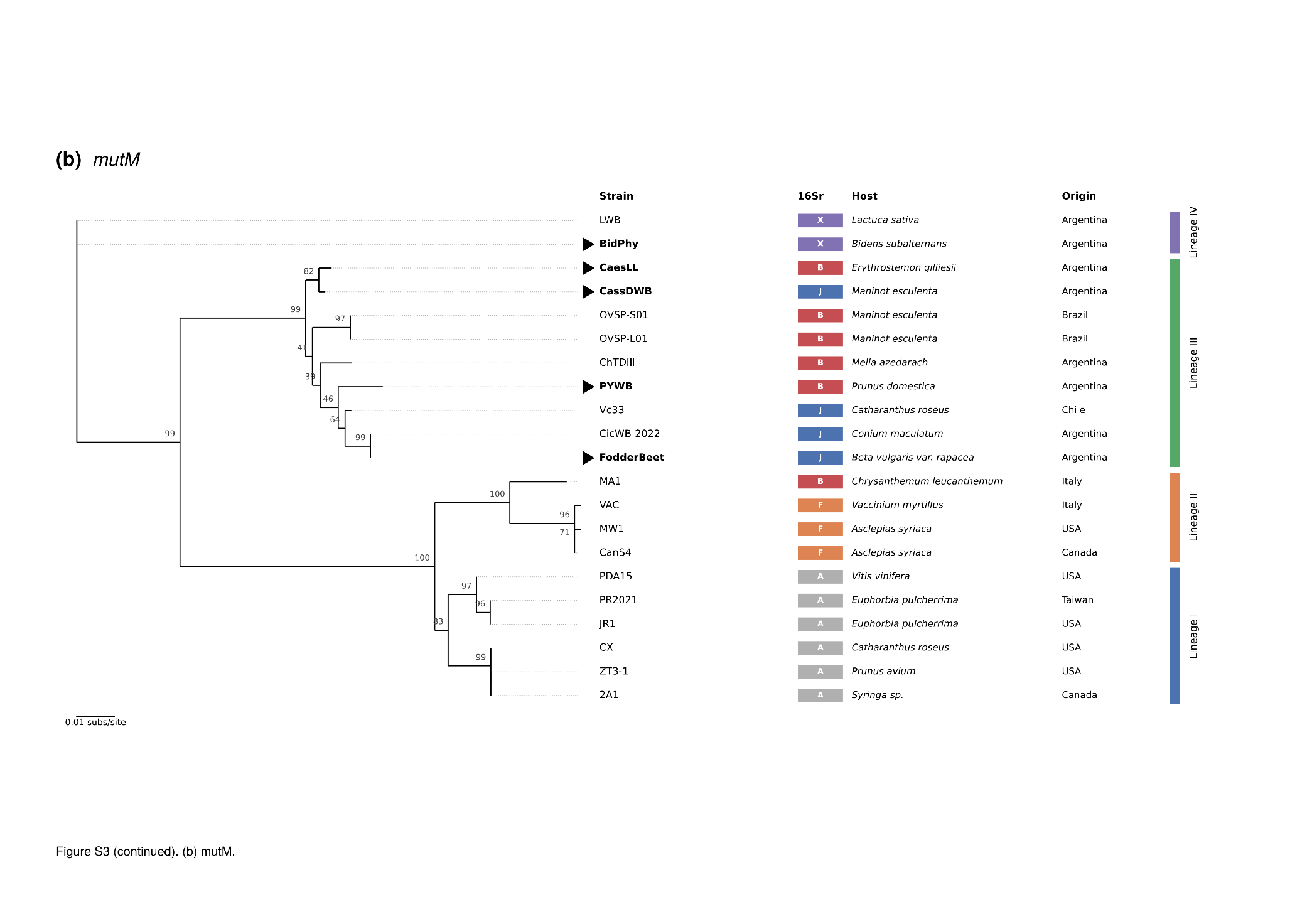

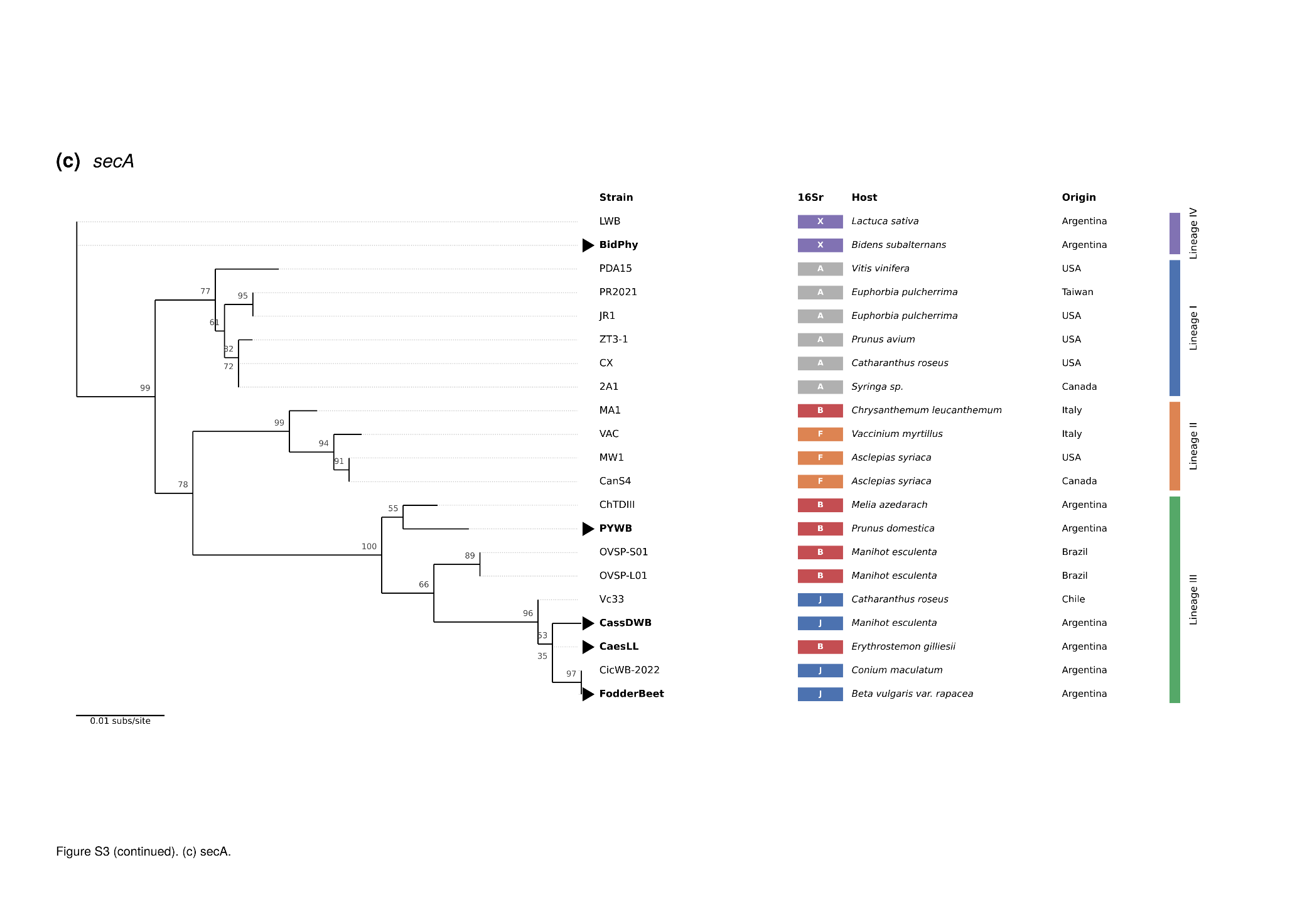

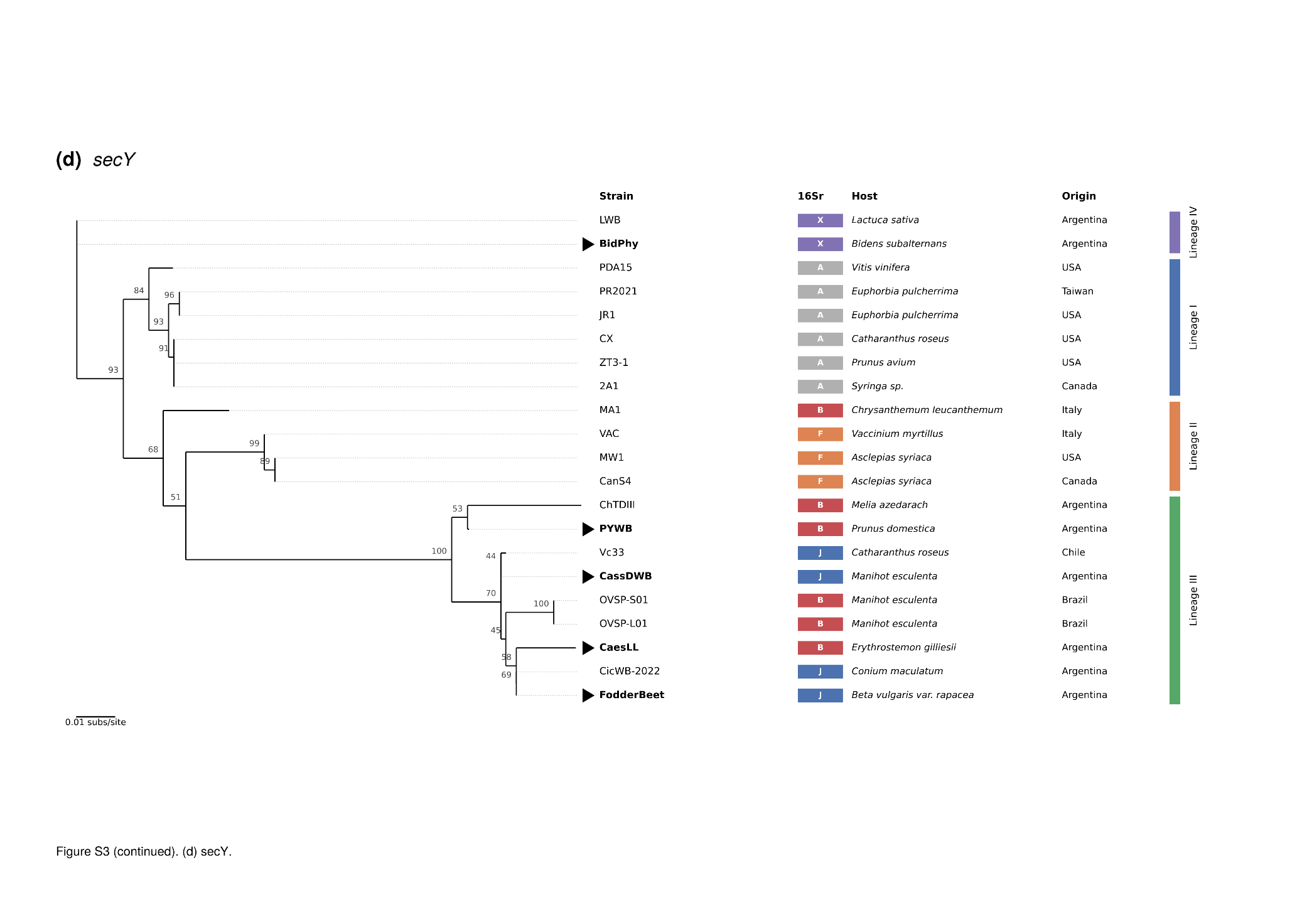

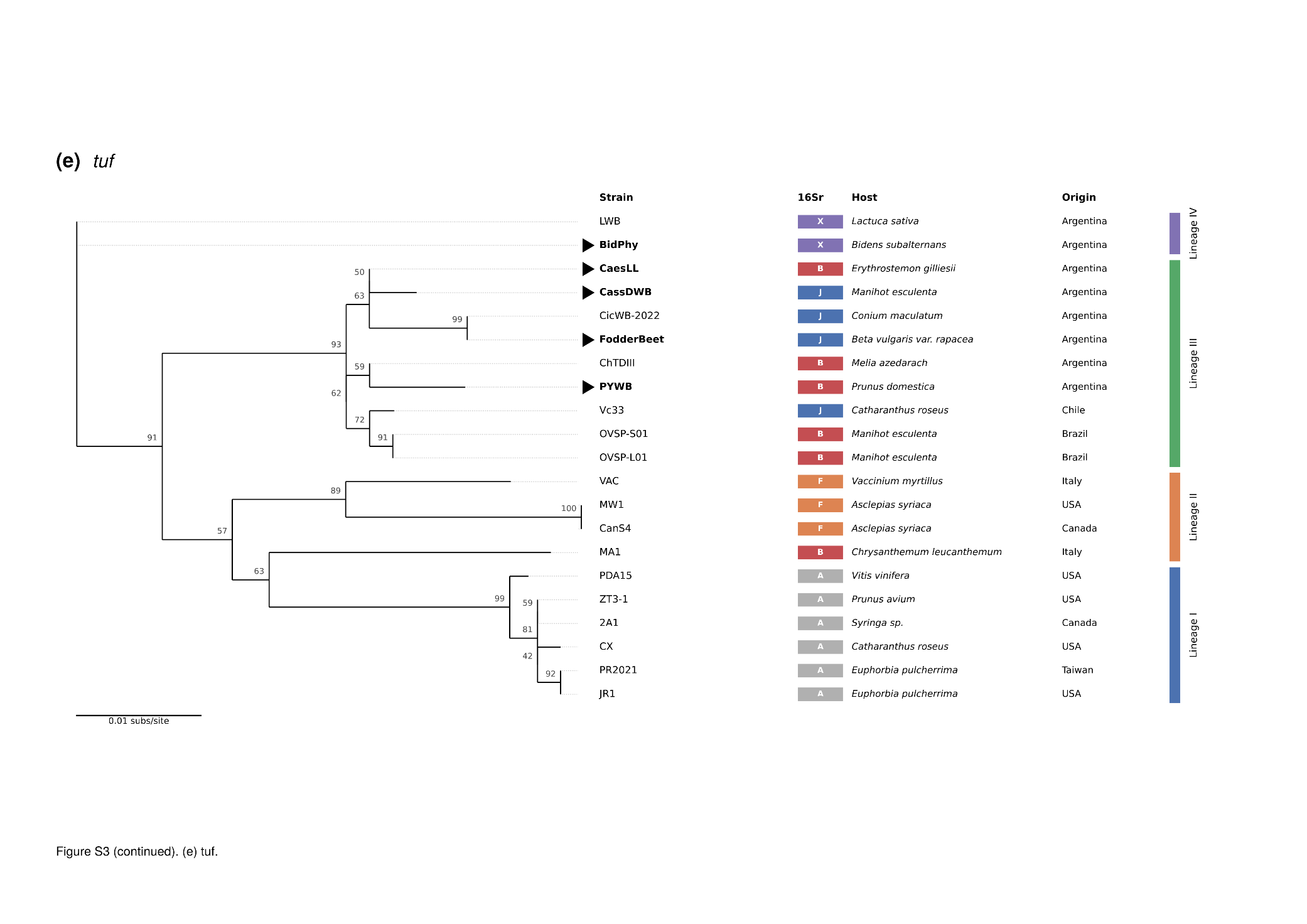

**Supplementary references (software)**

Hoang, D.T., Chernomor, O., von Haeseler, A., Minh, B.Q. & Vinh, L.S. (2018) UFBoot2: improving the ultrafast bootstrap approximation. *Molecular Biology and Evolution*, 35, 518–522.

Kalyaanamoorthy, S., Minh, B.Q., Wong, T.K.F., von Haeseler, A. & Jermiin, L.S. (2017) ModelFinder: fast model selection for accurate phylogenetic estimates. *Nature Methods*, 14, 587–589.

Manni, M., Berkeley, M.R., Seppey, M., Simão, F.A. & Zdobnov, E.M. (2021) BUSCO update. *Molecular Biology and Evolution*, 38, 4647–4654.

Marçais, G., Delcher, A.L., Phillippy, A.M., Coston, R., Salzberg, S.L. & Zimin, A. (2018) MUMmer4: a fast and versatile genome alignment system. *PLoS Computational Biology*, 14, e1005944.

Robinson, D.F. & Foulds, L.R. (1981) Comparison of phylogenetic trees. *Mathematical Biosciences*, 53, 131–147.

Shimoyama, Y. (2022) pyGenomeViz: a genome visualization Python package for comparative genomics. https://github.com/moshi4/pyGenomeViz
