## Supplementary material for "A genome-guided multilocus sequence analysis scheme to detect and resolve diversity within the X-disease phytoplasma group (16SrIII)": Table 1

**Table 1.** Genome assemblies of X-disease group (16SrIII) phytoplasmas analysed, outgroups and isolates sequenced in this study.

| **Strain** | **Lin.** | **16Sr** | **Host** | **Country** | **Accession** | **Level** | **Contigs** | **Size (kb)** | **G+C (%)** | **Compl. (%)** | **Ref.** |
| --- | --- | --- | --- | --- | --- | --- | --- | --- | --- | --- | --- |
| CX | I | III-A | *Catharanthus roseus*^e^ | USA | GCA_001277135.1 | Draft | 46 | 598.5 | 27.21 | 96.8 | 2 |
| 6A1 | I | III-A^b^ | *Syringa* sp. | Canada | GCA_033391615.1 | Draft | 2 | 625.2 | 27.32 | 97.1 | 12 |
| ZT3-1 | I | III-A^b^ | *Prunus avium* | USA | GCA_024586395.1 | Draft | 14 | 588.8 | 27.09 | 97.1 | 13 |
| PDA15 | I | III-A^b^ | *Vitis vinifera* | USA | GCA_041984945.1 | Draft | 61 | 595.0 | 26.96 | 95.2 | — |
| JR1 | I | III-A^c^ | *Euphorbia pulcherrima*^e^ | USA | GCA_000309465.1 | Draft | 185 | 631.4 | 27.31 | 95.2 | 1 |
| PR2021 | I | III-A^b,c^ | *Euphorbia pulcherrima* | Taiwan | GCA_029746895.1 | Complete | 2 | 709.9 | 26.65 | 97.1 | 5 |
| CanS4 | II | III-F^b^ | *Asclepias syriaca* | Canada | GCA_050286905.1 | Draft | 7 | 693.9 | 26.74 | 97.1 | 12 |
| MW1 | II | III-F | *Asclepias syriaca*^e^ | USA | GCA_000309485.1 | Draft | 158 | 583.8 | 27.43 | 95.2 | 1 |
| VAC | II | III-F | *Vaccinium myrtillus*^e^ | Italy | GCA_000309405.1 | Draft | 272 | 647.8 | 27.33 | 97.1 | 1 |
| MA1 | II | III-B | *Chrysanthemum leucanthemum*^e^ | Italy | GCA_000300695.1 | Draft | 197 | 597.2 | 27.13 | 91.4 | 1 |
| ChTDIII | III | III-B | *Melia azedarach* | Argentina | GCA_013391955.1 | Draft | 54^a^ | 733.2^a^ | 28.65^a^ | 97.1 | 4 |
| OVSP-S01 | III | III-B^b^ | *Manihot esculenta* | Brazil | GCA_059597155.1 | Draft | 1 | 599.9 | 27.82 | 97.1 | 8 |
| OVSP-L01 | III | III-B^b^ | *Manihot esculenta* | Brazil | GCA_059597165.1 | Draft | 1 | 601.5 | 27.79 | 97.1 | 8 |
| CicWB-2022 | III | III-J | *Conium maculatum* | Argentina | GCA_035853675.1 | Draft | 16 | 758.2 | 27.78 | 97.1 | 6 |
| Vc33 | III | III-J | *Catharanthus roseus*^e^ | Chile | GCA_001623385.2 | Draft | 29 | 687.1 | 27.87 | 86.4 | 3 |
| LWB | IV | III-X | *Lactuca sativa* | Argentina | GCA_051397825.1 | Complete | 1^f^ | 659.0^f^ | 27.03 | 97.1 | 7 |
| ‘*Ca.* P. phoenicium’ F1A | Outgroup | IX | *Prunus dulcis* | — | GCA_056945795.1 | Complete | 552.2 | 552.2 | 24.5 | 93.7 | — |
| ‘*Ca.* P. oryzae’ HN2022 | Outgroup | XI | *Oryza sativa* | China | GCA_027945135.1 | Complete | 511.3 | 511.3 | 20.5 | 91.5 | — |
| BidPhy | IV | III-X | *Bidens subalternans* | Argentina | MH497013^d^ | Sanger | — | — | — | — | 14 |
| FodderBeet | III | III-J | *Beta vulgaris* var. *rapacea* | Argentina | MH582072^d^ | Sanger | — | — | — | — | 15 |
| CassDWB | III | III-J | *Manihot esculenta* | Argentina | MF563363^d^ | Sanger | — | — | — | — | 9 |
| PYWB | III | III-B | *Prunus domestica* | Argentina | MT553102^d^ | Sanger | — | — | — | — | 10 |
| CaesLL | III | III-B | *Erythrostemon gilliesii* | Argentina | KC202812^d^ | Sanger | — | — | — | — | 11 |

^a^ After removal of 13 host-derived contigs (original assembly 67 contigs, 790.5 kb). ^b^ Subgroup assigned in this study by in silico 16S rRNA RFLP (F = 1.00) or by ANI ≥ 97% (ZT3-1, PDA15). ^c^ Variant of the 16SrIII-A pattern (F = 0.99); JR1 was originally reported as 16SrIII-H. ^d^ 16S rRNA accession of the isolate. ^e^ Strain maintained in the host indicated; see the original report for the plant of isolation. ^f^ Chromosome only; the plasmid pLWB-01 is not included. Completeness: CheckM, class *Mollicutes* marker set. Isolates (last five rows) were sequenced in this study for five MLSA loci (GenBank QB286830–QB286854; Table S3); —, not available or no associated publication. Lin: Lineage. References (genome or first report): 1, Saccardo et al. (2012); 2, Lee et al. (2015); 3, Zamorano and Fiore (2016); 4, Fernández et al. (2020c); 5, Pei et al. (2023); 6, Fernández et al. (2024); 7, Fernández et al. (2026); 8, Oliveira et al. (2026); 9, Fernández et al. (2018); 10, Bongiorno et al. (2020); 11, Alessio et al. (2025); 12, Pellegrinetti et al. (2026); 13, Wright and Harper (2022); 14, Fernández et al. (2020a); 15, Fernández et al. (2020b).
