## Supplementary material for "A genome-guided multilocus sequence analysis scheme to detect and resolve diversity within the X-disease phytoplasma group (16SrIII)": Table 2

**Table 2.** Primers designed in this study for the multilocus sequence analysis of X-disease group (16SrIII) phytoplasmas.

| **Locus** | **Primer** | **Sequence (5′→3′)** | **Ta (°C)** | **Amplicon (bp)** | ***In silico* (mismatches)** |
| --- | --- | --- | --- | --- | --- |
| *tuf* | tuf_Fw | GCTGCTCAAATGGATGCTGG | 62 | 598 | 0 in 16/16 |
|  | tuf_Rv | TAACTAAAACTTGTCCACGTTCTAC |  |  |  |
| *secA* | secA_Fw | CAGTTCATATCGTGACAGTTAATGA | 62 | 616 | R: 1 in OVSP (non-3′) |
|  | secA_Rv | CGACCATTTAAAATCCTACCAGTGA |  |  |  |
| *secY* | secY_Fw | TGAAAGAATGGCAAGAACAAGG | 63 | 729 | F: 1 in LWB (non-3′) |
|  | secY_Rv | CCCCTTCTAAGTAAGCATCTTGTT |  |  |  |
| *mutM* | mutM_Fw | TCTTAAGTGATGATTGGGTKCTTGT | 63 | 606 | F/R: 1 in OVSP; R: 1 at 3′ end in ZT3-1 |
|  | mutM_Rv | AAGAACCACGYCCGCCTATT |  |  |  |
| *cimH* | cimH_Fw | GCGCACATMATAACTGCTTTATT | 60 | 570 | F/R: 2 in LWB (non-3′) |
|  | cimH_Rv | TAATAACCCATTTTGTTTCCGAC |  |  |  |

Ta, annealing temperature. Degenerate positions follow IUPAC nomenclature (K = G/T; Y = C/T; M = A/C).

All reactions were run in 25 µL with Q5 High-Fidelity DNA polymerase: 98 °C for 30 s; 30 cycles of 98 °C for 10 s, Ta for 30 s and 72 °C for 30 s; and 72 °C for 2 min.
