## Supplementary material for "A genome-guided multilocus sequence analysis scheme to detect and resolve diversity within the X-disease phytoplasma group (16SrIII)": Table 3

**Table 3.** Characteristics of the MLSA loci (21 sequences: 16 genomes and five isolates; PHI: recombination test).

| **Locus** | **Length (nt)** | **Variable sites** | **Parsimony-informative** | **π** | **PHI p** |
| --- | --- | --- | --- | --- | --- |
| *tuf* | 530 | 53 | 42 | 0.029 | 0.55 |
| *secA* | 580 | 52 | 45 | 0.030 | 0.99 |
| *secY* | 695 | 95 | 82 | 0.053 | 0.18 |
| *mutM* | 566 | 99 | 91 | 0.068 | 0.77 |
| *cimH* | 499 | 113 | 98 | 0.074 | 0.14 |
| Concatenated | 2,870 | 412 | 358 | — | — |

Best-fit model for the concatenated alignment: TIM+F+G4 (MFP+MERGE merged the five loci into a single partition; BIC = 14,925.625)
