## Supplementary material for "A genome-guided multilocus sequence analysis scheme to detect and resolve diversity within the X-disease phytoplasma group (16SrIII)": Table 4

**Table 4.** Host range and geographic origin of the genomic lineages of the X-disease group (16SrIII), based on genome (SCG/ANI) or five-locus MLSA data.

| **Lineage** | **Strain / isolate** | **16Sr** | **Host** | **Host family** | **Host type** | **Country** | **Evidence** |
| --- | --- | --- | --- | --- | --- | --- | --- |
| I – ‘*Ca.* P. pruni’ *s.s.* | CX | III-A | *Catharanthus roseus*^a^ | Apocynaceae | Ornamental | USA | Genome |
|  | 6A1, ZT3-1 | III-A | *Prunus avium* | Rosaceae | Fruit crop | Canada, USA | Genome |
|  | PDA15 | III-A | *Vitis vinifera* | Vitaceae | Fruit crop | USA | Genome |
|  | JR1, PR2021 | III-A^b^ | *Euphorbia pulcherrima* | Euphorbiaceae | Ornamental | USA, Taiwan | Genome |
| II – 16SrIII-F/MA1 | MW1 | III-F | *Asclepias syriaca* | Apocynaceae | Wild plant | USA | Genome |
|  | VAC | III-F | *Vaccinium myrtillus* | Ericaceae | Wild plant / berry | Italy | Genome |
|  | CanS4 | III-F | *Asclepias syriaca* | Apocynaceae | Wild plant | Canada | Genome |
|  | MA1 | **III-B** | *Chrysanthemum leucanthemum*^a^ | Asteraceae | Wild plant | Italy | Genome |
| III – South American | ChTDIII | **III-B** | *Melia azedarach* | Meliaceae | Ornamental / forest tree | Argentina | Genome |
|  | OVSP-S01, OVSP-L01 | **III-B** | *Manihot esculenta* | Euphorbiaceae | Staple crop | Brazil | Genome |
|  | PYWB | **III-B** | *Prunus domestica* | Rosaceae | Fruit crop | Argentina | MLSA |
|  | CaesLL | **III-B** | *Erythrostemon gilliesii* | Fabaceae | Native shrub | Argentina | MLSA |
|  | CicWB-2022 | III-J | *Conium maculatum* | Apiaceae | Weed | Argentina | Genome |
|  | Vc33 | III-J | *Catharanthus roseus*^a^ | Apocynaceae | Ornamental | Chile | Genome |
|  | CassDWB | III-J | *Manihot esculenta* | Euphorbiaceae | Staple crop | Argentina | MLSA |
|  | FodderBeet | III-J | *Beta vulgaris* var. *rapacea* | Amaranthaceae | Forage crop | Argentina | MLSA |
| IV – 16SrIII-X | LWB | III-X | *Lactuca sativa* | Asteraceae | Vegetable crop | Argentina | Genome |
|  | BidPhy | III-X | *Bidens subalternans* | Asteraceae | Weed | Argentina | MLSA |

ᵃ Strain maintained in periwinkle (Catharanthus roseus) by graft transmission; for CX and Vc33 the host shown is the maintenance host, not the original one.

ᵇ Subgroup assigned in this study by *in silico* RFLP; the pattern was most like, but not identical with, the reference pattern of this subgroup (F < 1.00)
