## Supplementary material for "A genome-guided multilocus sequence analysis scheme to detect and resolve diversity within the X-disease phytoplasma group (16SrIII)": Table 5

**Table 5.** PCR amplification of MLSA loci from 16SrIII isolates and phytoplasmas of other groups (50 ng DNA; conventional PCR).

| **Isolate** | **Host** | **16Sr** | **16S** | **tuf** | **secA** | **secY** | **mutM** | **cimH** |
| --- | --- | --- | --- | --- | --- | --- | --- | --- |
| ChTDIII | *Melia azedarach* | III-B | + | + | + | + | + | + |
| CaesLL | *Erythrostemon gilliesii* | III-B | + | + | + | + | + | + |
| PYWB | *Prunus domestica* | III-B | + | + | + | + | + | + |
| BellVir | *Bellis perennis* | III-J | + | + | + | + | + | + |
| CassDWB | *Manihot esculenta* | III-J | + | + | + | + | + | + |
| FodderBeet | *Beta vulgaris* var. *rapacea* | III-J | + | + | + | + | + | + |
| BidPhy | *Bidens subalternans* | III-X | + | + | + | + | + | + |
| ArAWB | *Medicago sativa* | VII-C | + | – | – | – | – | – |
| ChTYXIII | *Melia azedarach* | XIII-G | + | – | – | – | – | – |
| BidPhy-Cba | *Bidens subalternans* | I-A/J | + | – | – | – | – | + |

+, amplicon of expected size; –, no amplification. 16S, R16F2n/R16R2 (positive control). ChTYXIII, ‘*Ca.* P. meliae’. BidPhy (isolate SJQ, 16SrIII-X) and BidPhy-Cba (16SrI) are different isolates from the same host species.
